# Transition scale-spaces: A computational theory for the discretized entorhinal cortex

**DOI:** 10.1101/543801

**Authors:** Nicolai Waniek

**Affiliations:** Bosch Center for Artificial Intelligence (BCAI), Robert Bosch GmbH, Robert-Bosch-Campus 1, 71272, Renningen, Germany

**Keywords:** grid cells, entorhinal cortex, spatial navigation, transition systems, scale-space, multi-resolution

## Abstract

Although hippocampal grid cells are thought to be crucial for spatial navigation, their computational purpose remains disputed. Recently, they were proposed to represent spatial transitions and to convey this knowledge downstream to place cells. However, a single scale of transitions is insufficient to plan long goal-directed sequences in behaviorally acceptable time.

Here, a scale-space data structure is suggested to optimally accelerate retrievals from transition systems, called Transition Scale-Space (TSS). Remaining exclusively on an algorithmic level, the scale increment is proved to be ideally 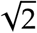 for biologically plausible receptive fields. It is then argued that temporal buffering is necessary to learn the scale-space online. Next, two modes for retrieval of sequences from the TSS are presented, namely top-down and bottom-up. The two modes are evaluated in symbolic simulations, i.e., without biologically plausible spiking neurons. Additionally, a TSS is used for short-cut discovery in a simulated Morris water maze. Finally, the presented results are discussed in depth with respect to biological plausibility, and several testable predictions derived. Moreover, relations to other grid cell models, multi-resolution path planning, and scale-space theory are highlighted. Summarized, reward-free transition encoding is shown here, in a theoretical model, to be compatible with the observed discretization along the dorso-ventral axis of medial Entorhinal Cortex (MEC).

Because the theoretical model generalizes beyond navigation, the TSS is suggested to be a general-purpose cortical data structure for fast retrieval of sequences and relational knowledge.

## 1 Introduction and Motivation

Spatial navigation and localization depend on a number of cells which represent spatial modalities (Rowland et al., 2016). Place cells, for instance, selectively respond to the location of an animal by firing only in distinct localized regions of an environment, called place fields (O’Keefe, 1979), and were found to be of critical importance for localization and path planning (de Lavilléon et al., 2015; Morris et al., 1982). The most perplexing cells are, however, grid cells of the Entorhinal Cortex (EC) (Hafting et al., 2005). Their eponymous periodic spatial firing patterns, called grid fields, regularly tessellate space. Embedded in an inhibitory local recurrent circuit (Couey et al., 2013), they interact with several other spatially modulated cells of the EC, for instance head direction (Ranck, 1984; Sargolini et al., 2006), border (Lever et al., 2009; Savelli et al., 2008), or speed cells (Hinman et al., 2016; Kropff et al., 2015). Moreover, they are influenced by the geometry of an animal’s surrounding (Krupic et al., 2014; Wernle et al., 2017), and project to place cells (Fyhn et al., 2008; E. I. Moser and M.-B. Moser, 2008). In turn, place cells recurrently project to the EC (Bonnevie et al., 2013).

Theoretical studies suggested early on that place fields can be generated by converging grid activity of different sizes (Franzius et al., 2007; Fuhs and Touretzky, 2006; Jeffery, 2007; McNaughton, Battaglia, et al., 2006; Solstad et al., 2006). The idea is that, while a single periodic grid response leads to ambiguous localization results, superimposed grid responses of several sizes cancel each other out and lead to unambiguous localized activity that is similar to place fields even in large environments. Afterwards it turned out that grid cells organize, in fact, in distinct modules along the dorso-ventral axis of the EC (Stensola et al., 2012): Cells within one grid module share the orientation of the hexagonal pattern, show a relative phase shift to each other, and express the same size with respect to their grid fields. Surprisingly though, the sizes of the grid fields increase from one module to the next in discrete steps with a scale factor that is suspiciously close to 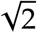.

Grid cells are considered a vehicle to understand higher cortical representations and functionality (E. I. Moser, Roudi, et al., 2014). This is not only due to their remarkable properties, but also because they are situated at the apex of the cortical processing hierarchy (Fellemann and Van Essen, 1991). They are commonly believed to perform either of two functions. On the one hand, hexagonal grid fields were shown to be suitable for path integration (Burak and Fiete, 2009; Fuhs and Touretzky, 2006; McNaughton, Battaglia, et al., 2006). On the other hand, they theoretically outperform place cells in localization tasks (Mathis et al., 2012; Stemmler et al., 2015), and were recently shown to emerge in recurrent models for vector-based navigation (Banino et al., 2018; Cueva and X.-X. Wei, 2018). However, the difference between most existing models lies primarily in the way that either of the two is achieved (see reviews in Giocomo et al., 2011; Shipston-Sharman et al., 2016; Zilli, 2012).

There are concerns about both perspectives. First, many path integration models quickly accumulate noise and require additional mechanisms to prevent drift (Burak, 2006; Burak and Fiete, 2009), especially when deployed in real-world scenarios (Mulas et al., 2016). It was also observed that place cell activity of preweaning rats depends on traveled distance, even though the grid cell system is still unstable during this stage of development (Bjerknes et al., 2018). Second, there is accumulating evidence that is in conflict with the idea that the grid code is a primary source for the formation of place cell activity and, thus, localization. For instance, grid cells emerge only after place cells during postnatal development (Langston et al., 2010; Wills et al., 2010). In addition, removal of projections from grid to place cells in lesion studies demonstrated that the place code is only partially influenced by the elimination of grid inputs (Chen et al., 2019; Hales et al., 2014). It also remains unclear how downstream neurons should read out the ambiguous hexagonal representation (Bush et al., 2014). Third, vector-based navigation models that use multiple scales of grid cells to represent a target vector typically generalize only up to an area that depends on the number of scales. That is, ambiguous target vectors will appear inevitably as soon as the surrounding environment with which the model has to cope is too large. Several grid coding strategies were proposed that are based on or related to residue number systems to address the issue of ambiguous responses (Fiete et al., 2008; Gorchetchnikov and Grossberg, 2007; Masson and Girard, 2010; Mathis et al., 2012; Sreenivasan and Fiete, 2011; X.-x. Wei et al., 2015). Although they significantly improve the issue, they still have a maximal distance after which the grid patterns repeat and are thus also subject to the third concern. Conclusively, the computational purpose of grid cells remains opaque and explanations of their peculiarities, namely their hexagonal grid fields as well as discrete scale increments, unsatisfying.

### 1.1 Contribution and organization

The main contribution of this paper is to analytically derive the optimal scale-increment of an acceleration data structure for retrievals of sequences from transition systems. More precisely, the computational problem that is considered is retrieval of goal-directed sequences from memory with behaviorally relevant performance. The results are based on prior research, which treated grid fields as individual elements of a larger structure (Waniek, 2018). Using the same approach, it is shown how acceleration of retrievals connects to existing discrete data structures and algorithms. Subsequently, this is extended to biologically plausible receptive fields and the optimal scale increment of 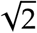 for this case proved analytically. In addition to the optimal scale increment, the results presented in this paper therefore also demonstrate how to relate cortical representations and computations to classical data structures and algorithms.

The paper is organized as follows. Subsection 1.2 introduces relevant nomenclature and concepts that will be used throughout this paper. For convenience, the section contains a summary in tabular form. Then, Section 2 presents related work. Subsequently, Section 3 presents the acceleration data structure and optimality results, which is split into the following parts. First, the necessity for acceleration is motivated using an example that considers neural dynamics. Second, a multi-scale data structure, called Transition Scale-Space (TSS), is presented that optimally accelerates retrievals given discrete representations of data. Third, bottom-up construction is shown for a TSS with neurally plausible receptive fields, which yields a scale-increment of 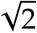 between consecutive scales of the TSS. Fourth, 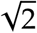 is proved analytically to be optimal for neurally plausible fields, and evaluated numerically. Fifth, it is argued that, for bottom-up construction, temporal buffering of data is required. In turn, this yields a relation between the theoretical model and Theta Phase Precession (TPP). Sixth, and closing this section, two modes for retrieval of sequences from the data structure are presented. After this, Section 4 presents simulations of the retrieval modes, as well as short-cut discovery capabilities in a simulated Morris water maze. Note that the experiments use symbolic simulations. Subsequently, Section 5 discusses both the theoretical and experimental results, gives detailed comparisons to related models, and highlights other related and influential research domains. Finally, Section 6 closes the paper with a brief summary and an outlook to future work.

This paper remains on an abstract algorithmic level, i.e., the second level of Marr’s suggested analysis of neural computations (Marr, 1982). It does not provide simulations of spiking neural networks that implement the algorithms and data structure. In addition, it omits input-output relations from and to other cortical areas when possible. Yet, this abstract perspective allows to derive several testable predictions for neural realizations that will be discussed in depth.

### 1.2 Nomenclature and concepts of Multi-Transition Systems

This paper builds on terminology, concepts, and mathematical symbols that were introduced in Waniek, 2018. For convenience, they are repeated here and the most important aspects briefly discussed. Table 1 summarizes the nomenclature and mathematical symbols that will be used throughout this paper. The table also provides an overview how the concepts of Multi-Transition Systems (MTS) are believed to relate to cells or networks of the Hippocampus. Further details about MTS, including formal proofs and an exhaustive discussion, can be found in Waniek, 2018.

**Table 1:** Nomenclature and neural interpretations. Note that indices will be dropped if clear from context.

| Symbol | Term | Correspondence or interpretation |
| --- | --- | --- |
| $\sigma_i$ | symbol | A place cell of CA1 |
| $\Sigma$ | symbolic space | All representations of place cells in CA1 |
| $\Gamma$ | spatial transition space | Representations formed by all grid cells |
| $\gamma_k$ | spatial transition bundle | Grid cell |
| $\gamma_{k,l}$ | spatial transition | Single grid response $l$ of a grid cell $k$ |
| $\delta_n$ | spatial symbol | Distributed representation of sensory modalities that uniquely identifies a location, e.g., via boundary vector cell activity |
| $\Delta$ | spatial input space | Entirety of spatial symbols |
| $M_\Sigma$ | Memory that implements $\Sigma$ | Place cells of CA1 |
| $M_\Gamma$ | Memory that implements $\Gamma$ | Grid cells of EC |

A transition bundle π {*τ*_1_ *, …, τ_N_* } is a set of *N* transitions *τ_k_*. Each transition is a tuple *τ_k_* (*σ_m_, σ_n_*) between symbols *σ_m_, σ_n_* ∈ Σ of an alphabet Σ. The *domain* of a transition contains the symbol from which a transition starts (also called source symbol), and the *image* of a transition contains the symbols to which it leads (also called target symbols). These terms are transitive to bundles, i.e., the domain (image) of a bundle is the set of symbols in which transitions of the bundle start (end). Note that, for instance in Reinforcement Learning (RL), transitions *τ_k_* are usually viewed as functions that map one symbol to another (Sutton and Barto, 1998).

Transition bundling is restricted by certain constraints. In particular, only those transitions with mutually exclusive source and target symbols can be bundled. In spaces where arbitrary transitions between any two symbols are possible, for instance in the case of temporal transitions between arbitrary symbols, a bundle contains only transitions that start at one source symbol. In complete metric spaces, bundling can be performed for all those transitions for which the intersection of domains and images is empty. For the two-dimensional case, Waniek, 2018 showed that bundling is maximized by a periodic hexagonal grid. In this case, transition bundles can be understood to form a dense packing of on-center and off-surround receptive fields, and the symmetry of this packing depends on the structure of the input space.

These concepts are thought to correspond to cells of the Hippocampal formation as follows. Place cells represent symbols *σ_i_* ∈ Σ in a neural associative memory *M*_Σ_, and receive spatially modulated, contextual, and possibly other cortical afferents. Of these, spatial inputs are believed to origin from a suitable sensory representation, such as head and boundary vector cell activity, denoted by Δ. Grid cells of a neural memory *M*_Γ_ learn spatial transitions directly on Δ, and convey this knowledge to *M*_Σ_. To achieve optimal transition bundling, each grid cell performs dense packing of on-center and off-surround receptive fields on Δ as part of its dendritic computation, depicted in Figure 1A. Each on-center is denoted as a spatial symbol *δ _j_* ∈ Δ, and dense packing of spatial symbols necessarily discretizes the input space due to a finite number of neurons and dendrites. Moreover, each spatial transition bundle associates to symbolic representations in *M*_Σ_ via co-activation learning. Thereby, a grid cell learns all feasible spatial transitions from place cells that are active at a certain spatial location to all place cells in the surrounding area, illustrated in Figure 1B. Summarized, the hippocampal-entorhinal loop is believed to form an MTS, illustrated in Figure 1C, and neurons of *M*_Σ_ can be understood to form nodes of a topological map, while transitions are edges between these nodes. Note that an MTS was proposed to store temporal transitions in a seprate memory *M*_Π_. However, this memory will be mostly omitted here and only mentioned when relevant.

**Figure 1:**
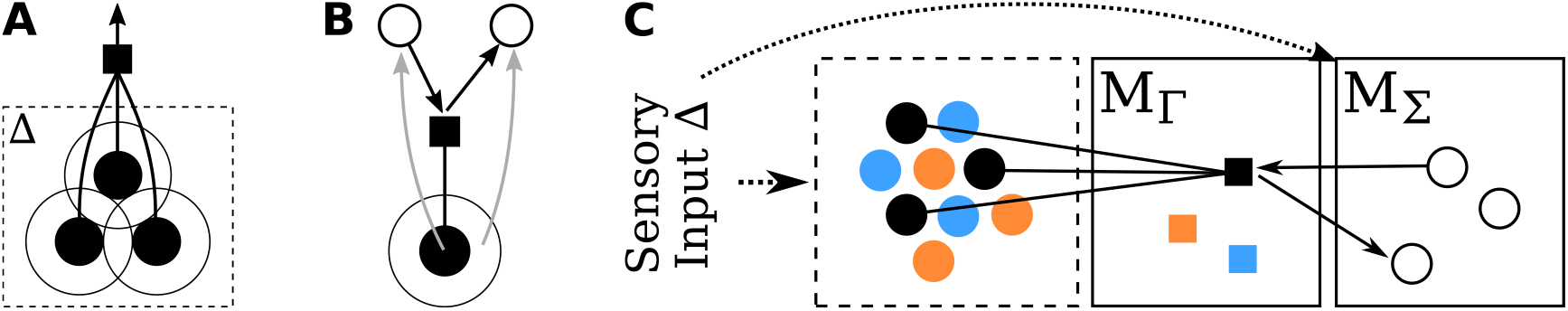
Conceptual overview of the hippocampal-entorhinal Multi-Transition System proposed in (Waniek, 2018). (A) A spatial transition bundle (black square) learns as many transitions as possible to maximize storage capacity of its dendritic tree (dense packing of on-center and off-surround circles). The input space on which this is performed is denoted Δ, and on-center areas are called spatial symbols *δ_i_* ∈ Δ. (B) A spatial transition bundle γ learns feasible transitions in a suitable input space and conveys this information to place cells *δ_i_, δ _j_* (white circles). (C) Learning a dense coverage of Δ requires multiple spatial transition bundles (grid cells; colored squares) in memory *M*_Γ_. These bundles convey spatial transition information to symbols *σ_i_* (place cells; white circles) of memory *M*_Σ_. Note that not all connections from transitions to bundles are depicted to improve clarity of the illustration.

Explicitly decoupling symbols (place cells) from spatial transition knowledge (grid cells) has several benefits. Most important, it renders place cells independent of the structure that is underlying sensory inputs. Consider an animal that wants to plan a path from its current location *A* to a remote target *B*, which means it needs to search a viable trajectory from *A* to *B*. This operation is called *expansion* of the path, denoted *A* ↝ *B*, into a sequence *A, σ*_1_ *, σ*_2_ *, …, B* of *N* ≥ 0 intermediate locations *σ_i_*. Because of the decoupling, an expansion needs to iterate only through symbol and transition memories, without any access to sensory representations. Also, a currently active representation of place cells may change abruptly, for instance due to a contextual change that induces a remapping event (Muller and Kubie, 1987). Relational spatial knowledge about feasible movements remains intact within grid cells during such an event, because grid cells are anchored to the sensory space independently from place cells as part of their dendritic computation. Also others recently noted the functional benefits of separating transitions that are recruited from sensory inputs from spatial representations that live independently of the structure of the sensory input (Whittington et al., 2018).

As a consequence of connectivity, place and grid cells were predicted to have independent but related recruitment mechanisms. Moreover, it was discussed that a network of such cells should express strong Winner-Take-All (WTA) dynamics to reduce overlap of symbols and, thereby, ambiguities in the representation. Couey et al., 2013 found that interactions between neurons in MEC are predominantly inhibitory, which is indicative of a strong WTA dynamics. Other recent evidence underpins the structure of network connectivity in an MTS and the independent recruitment processes for place and grid cells (Chen et al., 2019; Davoudi and Foster, 2019).

## 2 Related work

Soon after their discovery, several models were suggested that use multiple scales of grid cell responses to generate place cells (Fuhs and Touretzky, 2006; Jeffery, 2007; McNaughton, Battaglia, et al., 2006; Solstad et al., 2006). Others presented that multiple scales of grid like responses emerge when extracting slowly varying features from a spatially modulated input space and, likewise, used them downstream for localization purposes (Franzius et al., 2007; Wyss et al., 2006). Commonly, these models converge grid responses onto a single sheet of neurons, for instance using summation or multiplication. Models of this kind can be shown to yield unambiguous singly peaked activity similar to place codes even in environments that are significantly larger than the biggest grid period (Mathis et al., 2012). Moreover, X.-x. Wei et al., 2015 reported that – to reduce the number of required neurons for localization – grid fields ideally increase in discrete steps by a factor of 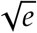, or about 1.4 − 1.7 when neurons fire stochastically. Others proposed that multiple scales of grid cells form a population vector for localization that can be read out using Bayesian inference (Stemmler et al., 2015), in which case the scale ratio was found to be ideally 3/2, or to improve representational capabilities by residue number systems (Fiete et al., 2008; Gorchetchnikov and Grossberg, 2007; Masson and Girard, 2010; Sreenivasan and Fiete, 2011). Although these values are in the range of measured scale increments, there exist doubts about the contribution of grid cells to localization (Bush et al., 2014). For instance, Hales et al., 2014 showed that the place code is only partially influenced when grid inputs were severed in lesion studies. More recently, Chen et al., 2019 presented evidence that self-motion and environmental cues have differential affects on place and grid cells, which contradicts the assumption that place cells are formed based on converging grid cell inputs. In contrast to these models, the work presented here is not based on localization, but on the idea that grid cells encode transitions or relational knowledge (Stachenfeld et al., 2016; Waniek, 2018). Moreover, while these models require multiple grid scales to solve ambiguities during localization, this paper argues that a discretized multi-scale representation is required to optimally accelerate retrievals from such transition knowledge.

The results presented here are conceptually related to vector-based navigation models, and in particular to the hierarchical linear look-ahead model presented in Erdem and Hasselmo, 2014 and Erdem, Milford, et al., 2015. It also shares several ideas with the navigation model by Redish and Touretzky, 1998, which predates the discovery of grid cells, and the linear look-ahead model by Kubie and Fenton, 2012. Brief summaries of these models, as well as detailed discussions of similarities and differences between these and the TSS model can be found in Subsection 5.3. Along the same lines of these models, Bush et al., 2014 argue that grid cells could potentially be used to compute linear target vectors between locations. They base their arguments on the observation that the entirety of grid fields of a single grid cell, itself discussed to be the result of path integration, might form a constant spatial metric. In turn, this could be used to compute translational vectors. However, Bush et al., 2014 do not present a mathematical or algorithmic model.

The paper assumes that knowledge in the Hippocampus is represented in the form of a relational graph or topological map, which was previously suggested in Dabaghian, Memoli, et al., 2012 and Dabaghian, Brandt, et al., 2014. It is also based on prior work that derived that a hexagonal distribution of fields is optimal to encode transitions in such a topological map (Waniek, 2018). It goes beyond this prior research by presenting how to optimally accelerate retrievals from such data.

Transition encoding in the Hippocampus was previously suggested and explored in a biologically plausible model (Cuperlier, Laroque, et al., 2004; Cuperlier, Quoy, et al., 2006). The model uses visual processing to extract landmarks, while transition cells support learning and planning of action sequences given visual stimuli. Moreover, it contains detailed biologically plausible hippocampal-prefrontal interactions. This model was evaluated on a robotics platform, and later extended via RL for the selection of an ideal trajectory to a target location (Hirel et al., 2010). Albeit influential for this paper, the authors neither addressed grid cells nor multiple scales of representation.

This work is related to the idea that grid cells perform Principal Component Analysis (Dordek et al., 2016), or that they compute a Successor Representation (Dayan, 1993; Momennejad et al., 2016; Stachenfeld et al., 2016). The relationship and differences between these and the model presented in this paper will be examined in detail in Subsection 5.3.

Gustafson and Daw, 2011 proposed a geodesic grid cell model. Their work was motivated by the idea that an animal needs to learn similarity between locally adjacent places and not across global distances for navigational purposes. In addition, the authors noted a similarity between spatial basis functions and the multi-scale representation of grid cells. Albeit the basic motivation for the geodesic grid cell model, MTS (Waniek, 2018), and the work presented here is equivalent, there are significant differences. For instance, while Gustafson and Daw, 2011 require a reward structure in the environment, both MTS and and the results presented here are derived independently of rewards. Moreover, while they chose predefined grid spacings, the optimal scale-increment for multi-scale representations will be derived analytically in Section 3. Further details on the relation between Gustafson and Daw, 2011 and Waniek, 2018 can be found in Appendix E.

Recently, McNamee et al., 2016 argued that neural networks that have to perform optimal planning operations should *in general* express discretized scales. Specifically, they proved that discretization minimizes the description length of a planning task and thereby achieves efficient encodings and optimal retrieval. However, their theoretical investigation did not link to grid cells.

The results presented here connect spatial navigation in the rodent brain with algorithms and data structures from robotics and computer science. For instance, Behnke, 2004 used representations with variable resolutions to accelerate robot navigation. Also, the presented results are intimately connected to scale-space theory, a formal mathematical framework that is widely used in the computer vision and signal processing communities (Lindeberg, 1994; Lindeberg, 2010; Witkin, 1983). These connections will be discussed in further depth in Subsection 5.4.

## 3 Scale-spaces for optimal transition look-ahead

This section examines the computational complexity of transition encoding during retrieval, and how this relates to behavioral necessities. Then, optimal acceleration of such retrievals using a scale-space structure is presented. Although provided in the context of spatial navigation, the results generalize to other transition systems. For instance, temporal transition systems for episodic memories are expected to form scale-space structures along appropriate dimensions.

Note that retrieval of sequences from a memory *M*_Σ_ requires activating a specific source symbol, and monitoring when a target symbol activates. This form of in- and output is expected to be the result of prefrontal-hippocampal interactions, similar to the prefrontral cortex columns and their simplifications to reward cells in the work by Erdem and Hasselmo, 2012 and, respectively, Erdem and Hasselmo, 2014. However, details about such interactions are unimportant for the scale-space structure and algorithms that are presented in this section, and are therefore omitted.

### 3.1 Computational complexity of retrievals and pre-play

Consider an MTS *ℳ* that is used for navigational purposes in an idealized two-dimensional input space Δ. *ℳ* consists of a finite number of symbols in a memory *M*_Σ_, and finite numbers of spatial and temporal transitions. The modular organization of *ℳ* is depicted in Figure 1C, and Figure 2B emphasizes the recurrent connectivity between the neural memories. Moreover, *ℳ* is subject to the following simplifications: (i) any uncertainty in the representation of the input space is ignored and (ii) spatial transitions organize according to a WTA mechanism. Then, spatial transitions perform a Voronoi tessellation that leads to hexagonal fields, illustrated in Figure 2A. These simplifications will be relaxed in Subsection 3.3. Finally, assume that the environment was explored sufficiently long and symbols and transitions were learned and stored in *ℳ* accordingly.

**Figure 2:**
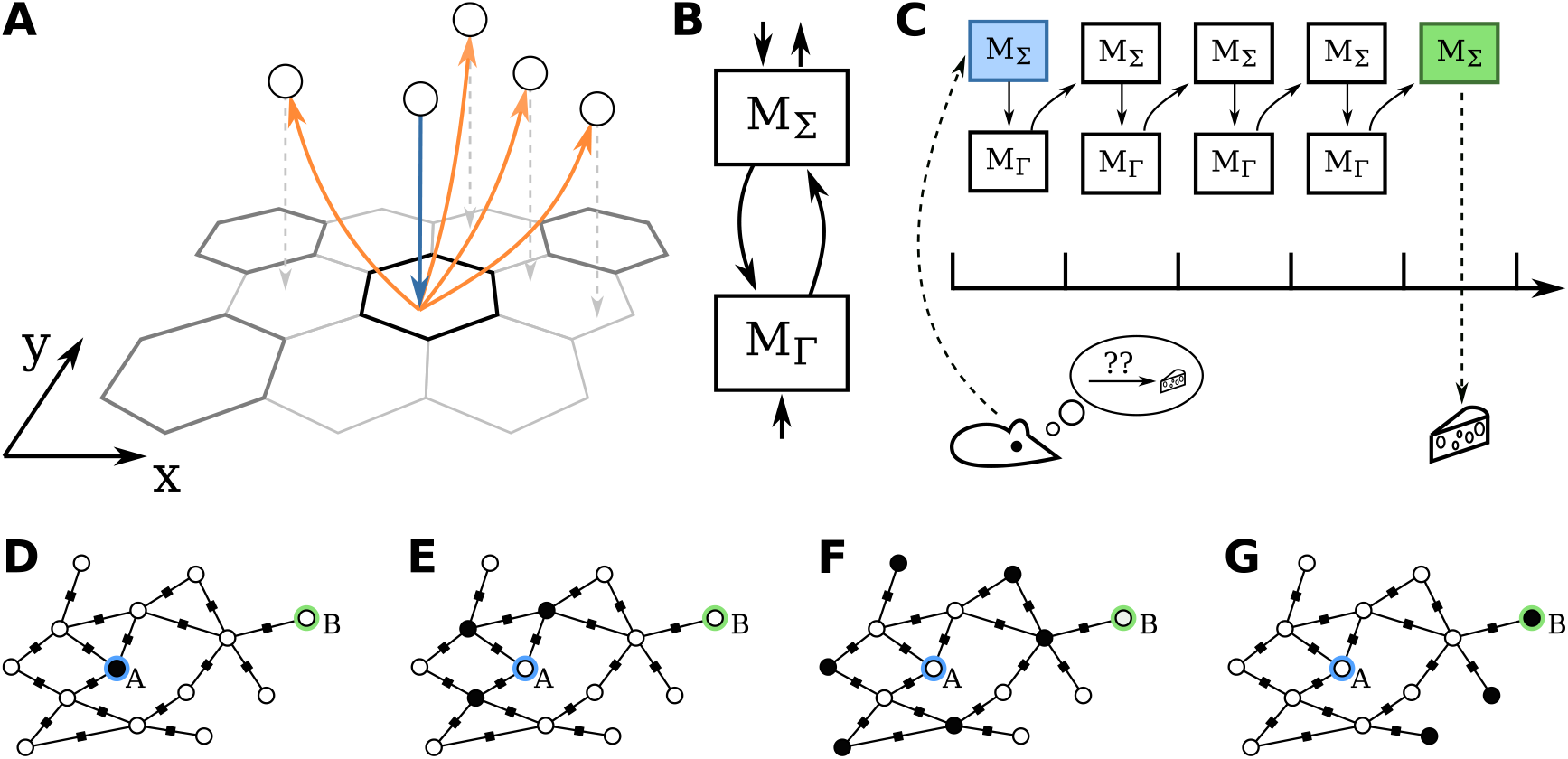
(A) Discrete Voronoi clustering on a suitable input space (indicated by the *x*-*y* axes) with absolute WTA dynamics yields hexagonal fields. A spatial transition (black hexagon) associates with all symbols that are co-active with its *domain* (circle with blue arrow) during exploration, and learns transitions (orange arrows) to symbols in its image (circles with gray dashed arrows). Dark gray, periodically repeating hexagons indicate the bundle to which the spatial transition belongs. (B) Memories of a MTS are recursively connected, shown only for *M*_Σ_ and *M*_Γ_. (C) Given activation of a starting symbol (indicated by blue color of *M*_Γ_) on a linear track that was discretized into bins (black arrow with vertical bars), mentally traveling to a target involves recursively activating *M*_Σ_ and the transition memories (only *M*_Γ_ shown here) until the target symbol is active in *M*_Σ_ (indicated by a green box). (D-G) Propagating wave of activity (black circles) in a graph when searching if a connected path exists between symbols *A* and *B*. Each edge involves a spatial transition, indicated by a small black box.

Determining the existence of a path between two symbols depends linearly on the number of recursive computations. In terms of *ℳ*, the path *A* ↝ *B* needs to be expanded into a sequence *A, …, σ_i_, …, B* of intermediary symbols *σ_i_*, which can be achieved as follows. First, the start symbol is activated in *M*_Σ_. Then, (indirect) recursive propagation of activity via transition memories will eventually activate *B*, if an expanded path from *A* to *B* exists. This simple algorithm can be depicted as a propagation algorithm on a graph because the data stored in an MTS forms a topological space (see Figure 2D-G). Moreover, the activity propagation is a flooding algorithm, which can be easily implemented in distributed processing systems (Raynal, 2013). The worst case scenario for this algorithm is when *A* and *B* are on the two ends of a linear track, in which case recursive retrievals via temporal transition memory *M*_Π_ and spatial transition memory *M*_Γ_ coincide. The runtime complexity resides in *O*(*N*), where *N* is the number of symbols on the shortest path between *A* and *B*.

The linear dependency on *N* is problematic especially when temporal dynamics of real neural networks are respected. Consider an implementation of *ℳ* that consists of neural associative memories, each with temporal dynamics that consume about 10 ms for spike generation, propagation, and axonal delays. Furthermore, let *A* and *B* be on a linear track with distance *D* 200 m and grid field size *d_g_* 20 cm. Hence, the expanded path from *A* to *B* consists of 1000 intermediate recursive evaluations of the *M*_Σ_ − *M*_Γ_ loop. This recursive retrieval is illustrated in a simplified manner in Figure 2C. Overall, recursive path expansion consumes 20 s to mentally travel from *A* to *B*. The runtime is expected to increase further when the animal has to choose from a set of candidate solutions, for instance with some reward propagation mechanism or graph search, and especially with stochastic neural responses. Moreover, the expansion must be repeated for each novel target. This behaviorally questionable performance for model-based algorithms was previously noted in the context of grid cells by Dordek et al., 2016 and Stachenfeld et al., 2016. Summarizing, recursive retrieval in a single-scale transition system has a time complexity that is impractical from a behavioral perspective.

Another problem is combinatorial complexity. Recall that transition systems induce a graph, in which vertices correspond to states or symbols, and edges to transitions between symbols. The number of vertices and edges in such a graph grows exponentially with the number of dimensions of the input space. Moreover, MTS assume constant cost to move from one symbol to another, which means that the weights of all edges are constant. Thereby, the number of feasible paths between two vertices, but also the actual runtime for shortest-path computations, grows similarly. In principle and given the assumption that each vertex of the graph is an independent computational unit, such shortest path computations can be parallelized easily using flooding algorithms (Lynch, 1996; Raynal, 2013). Then, shortest-path computation corresponds to a traveling wave of messages from source to target. Although this reduces searching to the linear case that was described above, it is still affected by the aforementioned problematic runtime complexity.

### 3.2 Optimal retrievals in discrete transition systems

Consider a spatial transition *γ_i_* of an MTS with discrete representations. Then, *γ_i_* informs about transitions jointly in spaces Δ and Σ, depicted in Figure 3A. The data represented by *γ_i_* forms two conjoint linked-lists, illustrated in a simplified manner in Figure 3B. Then, acceleration of retrievals can be performed optimally by an *interval skip list* (see Appendix A for a brief introduction to this data structure and how it relates to binary search).

**Figure 3:**
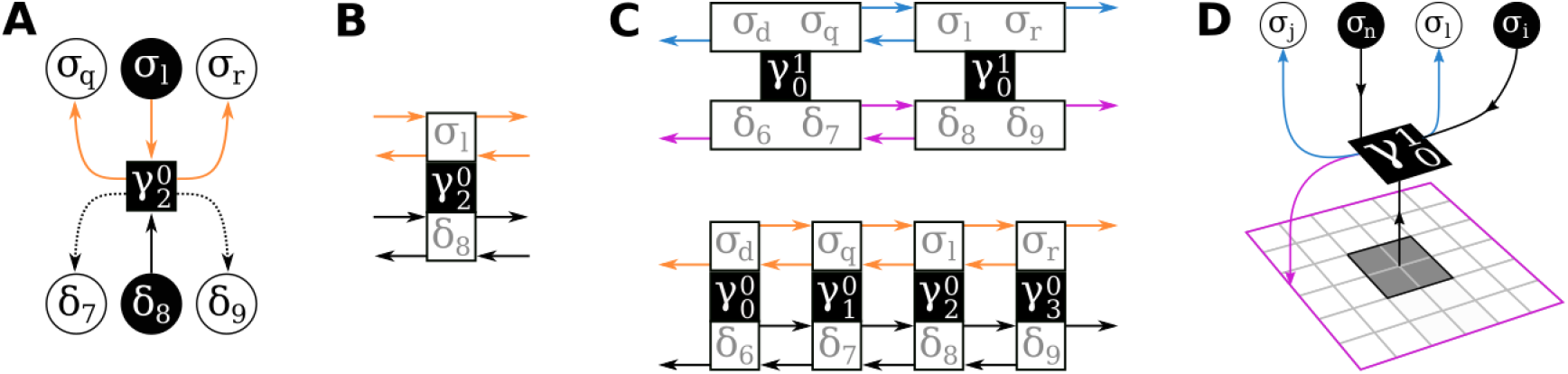
Scale-space construction and co-activation learning in discrete spaces. (A) Recall that a spatial transition represents transitions in Δ and, simultaneously, in Σ (cf. Figure 1). While spatial symbols *δ_i_* ∈ Δ need to be spatially neighboring (consecutive indices), symbols *σ_j_* ∈ Σ need to be temporally adjacent. In a neural realization, solid arrows are hypothesized to be axonal connections, dashed arrows could be the result of on-center and on-surround dynamics. (B) Each transition *γ_k_* ∈ Γ can be depicted as an element of two conjoint interval skip lists (cf. Figure 13). (C) The domains of transitions on scale 0 of a discrete Transition Scale-Space 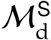 contain singular elements of the corresponding input spaces (bottom row). For optimality, transitions on a scale *s >* 0 merge the domains of two transitions of scale *s* − 1 (top row, shown for scale *s* = 1). (D) Two dimensional example that emphasizes the on-center (entire gray shaded area, bottom row) and off-surround (entire area enclosed by magenta border, excluding gray shaded region, bottom row) receptive field of a transition (black box) and co-activation learning symbols *σ* (top row: co-active symbols shaded black, transition leads other symbols via blue arrows). The gray grid is depicted for illustrative purposes only and to highlight the increase of the size of domain and image (cf. interval skip list, Figure 13B).

The construction procedure for interval skip lists directly translates to an optimal discrete Transition Scale-Space 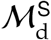. Let scale 0 be the lowest level of 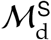. On scale 0, the domain of a spatial transition 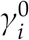 corresponds to a spatial symbol *δ_i_*. Its image is given by symbols surrounding *δ_i_*, for instance *δ_i_*_−1_ and *δ_i_*_+1_ in a one-dimensional setting, illustrated in Figure 3C. The domain of a spatial transition 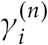 on scale *n >* 0 corresponds to the integrated domains of 2 adjacent transitions of the previous scale along each dimension, where 2 follows from constructing an optimal search data structure (de Berg et al., 1997; Knuth, 1998; Pugh, 1990). Likewise, the image of 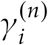 grows by a factor of 2 along each dimension with respect to scale *n* − 1 and encloses all symbols that surround the domain of 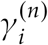. Note that the construction process is independent for each scale of the hierarchy. That is, each scale can be constructed without interactions from other scales when random access to afferents from Δ and Σ is provided. Panel D of Figure 3 illustrates the on-center and off-surround properties of a transition for the case of scale *s >* 0 in two-dimensions. Importantly, a transition on this scale leads to all symbols in its image simultaneously. An illustration that further simplifies the depiction of transitions is presented in Panel A of Figure 5. The panel plots only on-centers of receptive fields of transitions *γ*, and omits all arrows from transitions to symbol spaces.

### 3.3 Bottom up construction with biologically plausible representations

The previous section presented optimal acceleration of sequence retrievals for discrete data. Specifically, the set of spatial transitions in a discrete TSS forms two conjoint interval-skip lists, one operating on elements of Δ, the other on Σ (Figure 3C). In the discrete case, classical optimality results require that the intervals in such data structures double in size from one level of the hierarchy to the next. However, responses of neurons typically follow bell-shaped tuning curves (Butts and Goldman, 2006; Jazayeri and Movshon, 2006). Hence, this section will address biologically plausible representations and how to construct the data structure bottom-up with them.

Recall that an MTS bundles transitions to minimize the number of required neurons. To this end, spatial transition bundles densely pack on-center and off-surround receptive fields, where each center-surround field corresponds to a feasible spatial transition from one location to its surrounding region (see Subsection 1.2 or Waniek, 2018). Of particular importance is that each center-surround field is treated as an individual part of a bundle, depicted in Figure 4A. Consequently, the following will examine a single receptive field in isolation.

**Figure 4:**
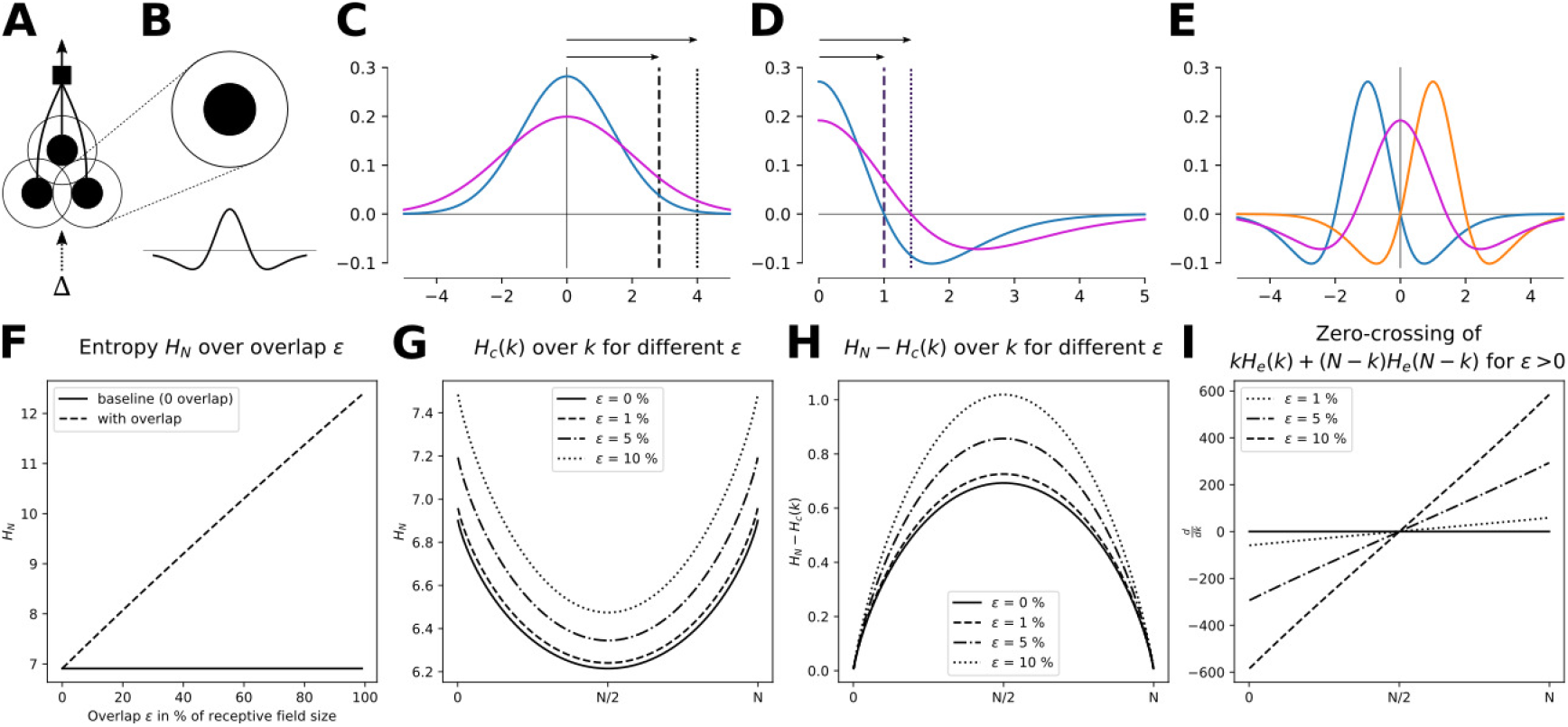
(A) Each spatial transition bundle receives afferents from a spatially modulated input space Δ and densely packs on-center and off-surround receptive fields. (B) Each receptive field can be treated independently and is modelled as Difference of Gaussians (DoG). (C) Bottom-up construction of the scale-space structure doubles the variance 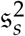 of the on-center Gaussian. This scales the width by 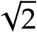, marked here for the 2𝔰*_s_* boundary (dashed and dotted lines). (D) Scaling the standard-deviation of a DoG by 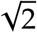 also scales the roots by 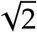, shown here for the positive domain of a DoG with 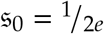 (blue) and 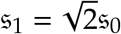 (magenta). (E) Convolution of two consecutive DoGs *X* and *Y* (blue, orange), parameterized by 𝔰_0_ and *µ_X_* −*µ_Y_*, leads to a DoG that is parameterized by *µ_Z_* 0 and 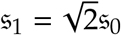 (magenta). (F-I) Total, cut, difference, and derivative of entropy in binary search for overlapping receptive fields (*N* = 1000 in panels G-H; see Subsection 3.4 for more details).

**Figure 5:**
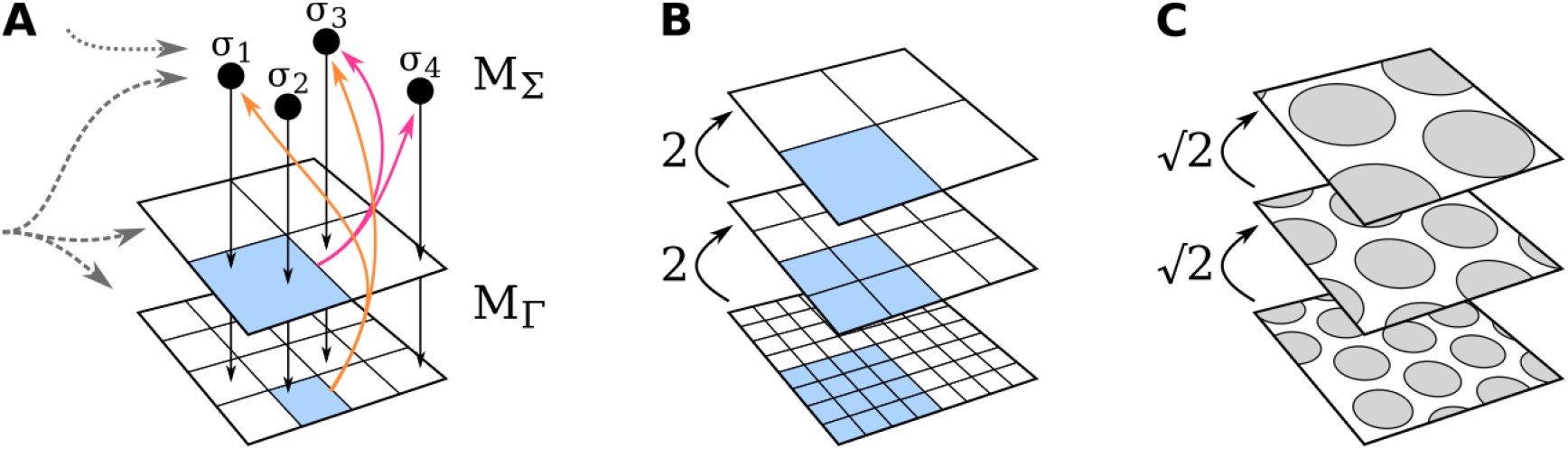
(A) Simplified illustration of learning transitions in a scale-space. Inputs to the Transition Scale-Space 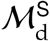 are indicated as gray dashed arrows. Transitions of a memory *M*_Γ_ learn their domains on the input space according to their scale (blue boxes), and relay this structural information to symbols *σ* ∈ Σ of memory *M*_Σ_ (black dots). In this example, a transition that is defined for *σ*_2_ leads to *σ*_1_ and *σ*_3_ on the lowest scale (orange arrows), whereas it leads to *σ*_3_ and *σ*_4_ on scale 1 (magenta arrows). (B) In the discrete case, the optimal scale increment of the domain (receptive field) of transitions is 2 along each dimension. In two-dimensions, illustrated here, this leads to the well known quad-tree data structure. (C) In stochastic input spaces that can be described with normally distributed pdfs, the scale increment from one scale to the next is 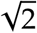.

Moreover, each receptive field is decomposed into separate center and surround components. This is motivated from data and models for retinal ganglion cells (Koenderink and van Doorn, 1990; Rodieck, 1965). In these models, field components were integrated either linearly (Marr and Hildreth, 1980; Rodieck, 1965), or with other mechanisms (Furman, 1965; Grossberg, 1970), all of which can be combined into one formal framework (Neumann et al., 1999).

Following these prior models, each center-surround receptive field is modelled as a Difference-of-Gaussians (DoG), illustrated in Figure 4A-B. Specifically, let *f* (*x*; *δ _j_*) denote the response function of a receptive field, parametrized by its preferred stimulus *δ_j_*, and given by

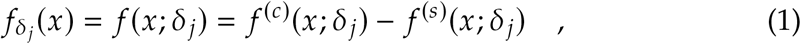

where *f* ^(*c*)^ and *f* ^(*s*)^ are the center and surround, respectively. To simplify the analysis, *f* ^(*c*)^ and *f* ^(*s*)^ are assumed to be isotropic Gaussians, e.g.

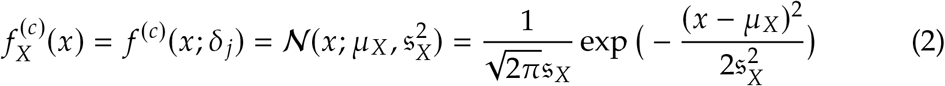

where *X* : *δ _j_* denotes the preferred stimulus, 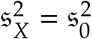 is the variance and thus width of the field, and the mean *µ_X_* : *µ*(*δ _j_*) depends on the preferred stimulus *δ _j_*. Note that all transitions necessarily adhere to the Cramér-Rao bound, which presents a lower bound with which a decoding scheme can extract information from pre-synaptic inputs (Cover and Thomas, 2006; Pilarski and Pokora, 2015). That is, *f* ^(*c*)^ of a spatial transition on the smallest scale *s* = 0 of transitions determines a highest possible resolution with which elements *δ* ∈ Δ can be distinguished. It is therefore sufficient to consider the size of *f* ^(*c*)^ on scale 0 to derive receptive field sizes on higher scales.

As described above, the constructive process to build an additional scale in an interval segment tree integrates two intervals of the previous scale. Moreover, this new interval is centered between the two smaller intervals *by construction*. In terms of receptive fields, the on-center region on a scale *s >* 0 needs to merge the representations of two receptive fields of the previous scale, which is clearly described by a convolution (Seeger and Volle, 1995). Hence, 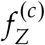(*x*) for a spatial transition on scale *s* 1 can be derived using the convolutional theorem as follows.

Let *ℱ* be the Fourier transform, and *ℱ*^−1^ its inverse, and let 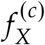 and 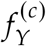 be the on-center components of two consecutive receptive fields on scale 0. That is, they are given by

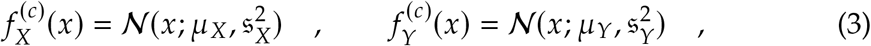

where 𝔰_X_ = 𝔰_Y_ = 𝔰_0_. Then (see Appendix B),

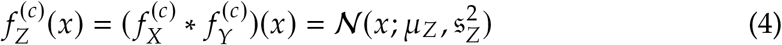

where

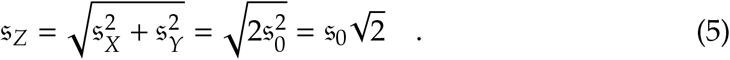

Furthermore, *µ_Z_ µ_X_* + *µ_Y_*. By construction, *µ_X_* −*µ_Y_* ⇒ *µ_Z_* 0.

The above results yield the following characteristic properties of receptive fields on higher scales, and of transition bundles to which they belong.

First, assume that the receptive field response is determined only by the on-center region of fields that are modelled as DoG. Then, the response is characterized by a Probability Density Function (pdf) in the form of a Gaussian. On the smallest scale *s* 0 of the data structure, this pdf is parameterized by variance 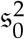 and the mean follows from the preferred stimulus. According to Equation 5, the variance on scale *s* = 1 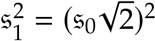. Recursively applied for higher scales, this yields

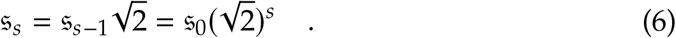

Because the Gaussians were assumed to be isotropic, this result generalizes directly to higher dimensions. That is, the covariance matrix 𝔖_0_ of the multivariate Gaussian is a diagonal matrix of the form 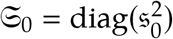 and higher scales follow according to Equation 6. Intuitively, the result means that the integration area of the receptive field, determined by the variance of the representation, doubles. Finally, assume that there is only a response if the input stimulus falls within *l*𝔰*_s_, l >* 0*, l* ∈ ℝ, for instance for *l* 2 → 2𝔰*_s_*. By Equation 6, the width (or radius) of the receptive field also increases by 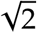 from one scale to the next. This property is illustrated in Figure 4B.

Now, assume that the response on scale *s* is determined jointly by on-center and off-surround dynamics of a DoG. Then, the field is defined by 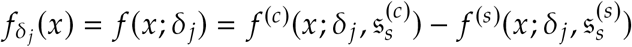. Because both *f* ^(*c*)^ and *f* ^(*s*)^ are Gaussians, they relate to each other by 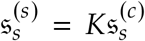 with *K >* 1. To improve terseness, the super-script ^(*c*)^ will be dropped in the following, i.e., 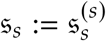. It is straightforward to analytically derive that a (one-dimensional) DoG has roots at

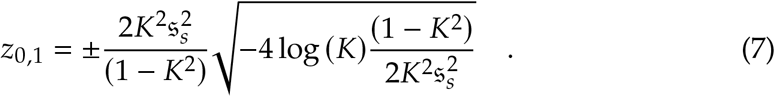

From this, it follows immediately that *r*𝔰*_s_* ⇒ *rz*_0,1_. That is, scaling the standard deviation 𝔰*_s_* by a factor *r* ∈ ℝ_+_ scales the roots also by *r*, irrespective of the value of *K*. An example of this is depicted in Figure 4D. Note that this result also generalizes to higher dimensions, again because of the assumed isotropy of the Gaussians. To conclude, the width (or radius) of the response increases according to the scaling factor *r*. Following the results from above, constructing the data structure bottom-up by convolution yields *r =* 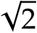, and thus the on-center region of receptive fields increase by 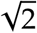, illustrated in Figure 4E.

Figure 5B and C depict simplified illustrations of discrete and a probabilistic scale-spaces that are constructed by this process.

Finally, these properties are expected to also increase the grid period by 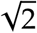. One reason is that in an optimal interval skip list, a transition leads to neighboring intervals that have the same size (cf. Figure 13B). Another reason is that in an MTS, each (spatial) bundle densely packs transitions to maximize dendritic capacity (cf. Subsection 1.2). Given the off-surround area that is required to avoid violation of logical considerations of transition bundling, dense coverage of an entire input space requires multiple bundles. Following arguments in Waniek, 2018, an entire network of bundles is expected to reduce overlap of on-center regions by competition to avoid ambiguities. Hence, these competitive interactions of transition bundles are conjectured to generate a push-pull mechanism that leads to grid periods that depend on 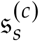 and increases the grid period consecutively by 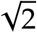. Intuitively, while a single transition bundle tries to densely pack receptive fields, competing transition bundles push them apart to insert their own receptive fields. Conclusively, this generates a dense coverage of the spatial input space Δ with on-center receptive fields of the same size, and a grid period that also follows from the size of on-center areas.

### 3.4 On the optimality of the 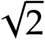 scale increment

The previous section presented that bottom-up construction of a Transition Scale-Space (TSS) with biologically plausible receptive fields yields a scale increment of 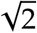. This will now be shown to be optimal using tools from information theory.

Recall the duality between interval skip lists and binary search (see Appendix A). Following results from Waniek, 2018, suppose that an MTS optimally covers a spatial input space with a finite number of spatial transitions. That is, they are uniformly distributed on and discretize the input space into spatial symbols *δ_i_, i* = 1*, …, N*. Conclusively, the necessary conditions for duality are met and it is sufficient to demonstrate optimality of the representation for one-dimensional binary search. As above, receptive fields are assumed to follow isotropic Gaussian tuning curves and can be treated in isolation.

Let *x* ∈ *X* be an input stimulus of a spatially modulated input space *X*. Moreover, let *f* (*x*; *δ_i_*) denote the on-center region of a receptive field of a single transition. Then, the indicator function

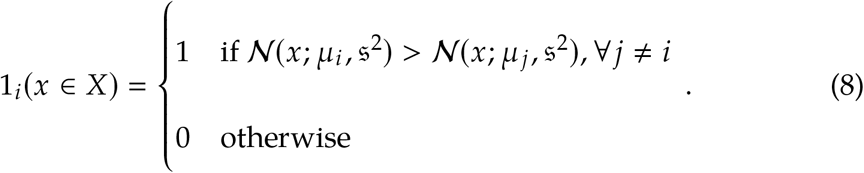

models WTA dynamics. That is, the function is 1 when an input stimulus *x* is closest to *δ_i_*.

In general, the probability for *f* (*x*; *δ_i_*) to win is thus

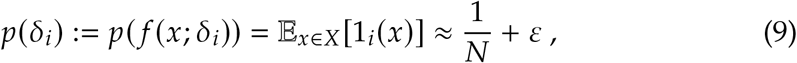

with 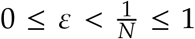. Here, *ε* accounts for overlapping receptive fields and to model stochasticity in WTA dynamics, in the following called called *overlap* for brevity. Note that 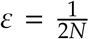 means a 50% overlap of two neighboring receptive fields.

The Shannon entropy *H_N_* (*X*) of a system of *N* such receptive fields is then

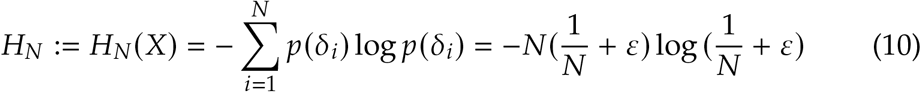

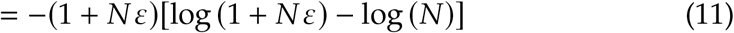

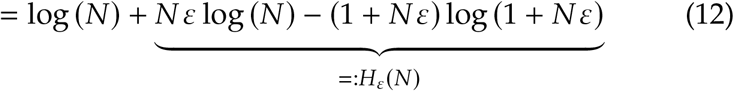

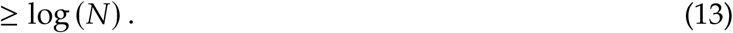

The entropy is lower-bounded by log (*N*) because *H_ε_*(*N*) *>* 0 for *ε >* 0 and lim*_ε_*_→0_ *H_ε_*(*N*) = 0. Panel F of Figure 4 illustrates how the entropy increases with increasing *ε* in a system of *N* = 1000 receptive fields. Also note that

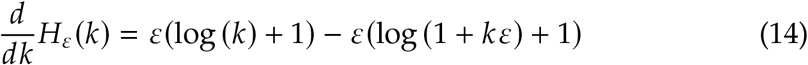

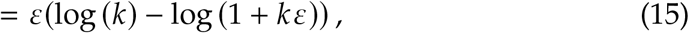

and that 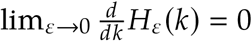.

Binary search splits the representational space at the *k*-th element, with *k* receptive fields to the left of the split, and *N* − *k* to the right. The entropy *H_c_* (*k*) of a split (or cut) of *N* receptive fields at the *k*-th item is

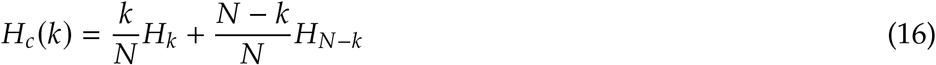

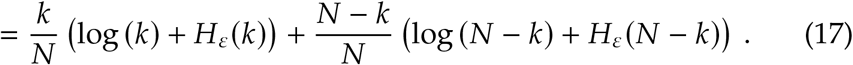

Panel G of Figure 4 depicts *H_c_* (*k*) for different overlaps *ε*.

The optimal split can be found by maximizing *H_N_* − *H_c_* (*k*). Specifically,

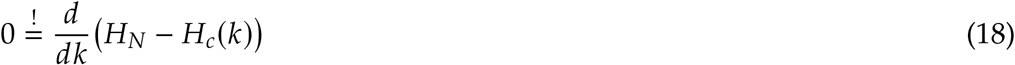

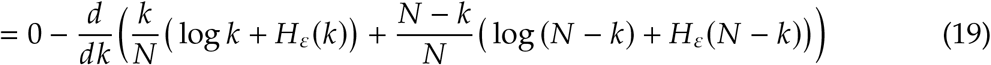

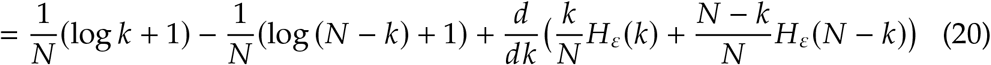

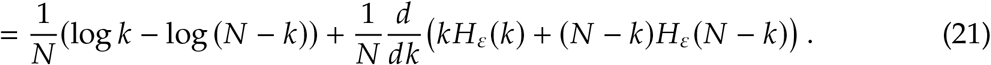

Numerical computations of *H_N_* − *H_c_* (*k*) are illustrated in panel H of Figure 4.

First consider *ε* = 0. In this case, Equation 21 reduces to the standard result for binary search, i.e.

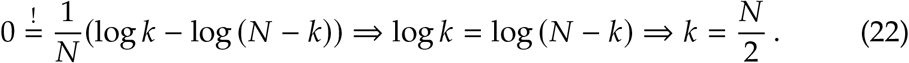

Now, assume that *ε* ≠ 0. Then, Equation 21 also holds for 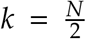. This is because *H_ε_*(*k*) is monotonically increasing and because of the symmetry of *kH_ε_*(*k*) + (*N* − *k*)*H_ε_*(*N* − *k*) around zero. Panel I of Figure 4 shows numerical computations of Equation 21 for *ε >* 0. Thus

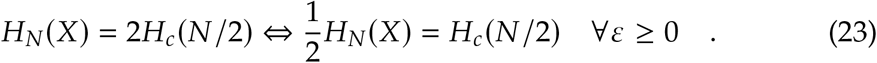

In other words, the uncertainty that is modelled by *ε* does not shift the optimality result for binary search and, therefore, consecutive steps of binary search ideally bisect the remaining entropy.

Now, consider the (differential) entropy 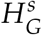 of a Gaussian function *G* with variance 𝔰^2^ at step *s* of the binary search, or in other words on level *s* of the TSS. Using Equation 23 it follows immediately that

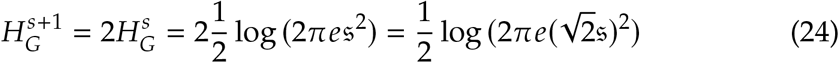

is optimal for level *s* + 1.

Conclusively, scaling the standard deviation by 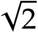 is optimal when building a TSS with Gaussian tuning curves. In addition, the results reveal that, on each scale, overlap of receptive fields and uncertainty in WTA dynamics, modelled by *ε*, should be minimized. Note that the results presented in this section hold for any data structure with Gaussian tuning curves that is dual to binary search.

### 3.5 Temporal integration, buffering, and bottom-up learning

The previous sections introduced the Transition Scale-Space (TSS) and derived the increment of the spatial receptive field analytically. Given random access to symbols *σ_i_* ∈ Σ, the data structure can be computed offline. It is, however, unlikely that an animal has this random access and must learn the data structure bottom up. Hence, this section examines how the temporal integration window changes between scales, and how on-line learning can be implemented.

Any *γ* ∈ Γ represents transitions in the spatial input space Δ and the symbolic space Σ simultaneously. Figure 3 illustrates this as two conjoint interval skip lists. That means that each *γ* ∈ Γ has a spatial domain dom_Δ_ and image im_Δ_ in Δ, but also a domain dom_Σ_ and image im_Σ_ in Σ. The previous sections showed that, to learn a transition *γ*^(*s*)^ on a scale *s >* 0, *γ*^(*s*)^ needs to ingrate multiple spatial symbols of Δ. Certainly, this also applies to afferents from Σ.

Learning a transition *γ*^(*s*)^ on scale *s >* 0 requires to integrate multiple symbols from Σ. However, recall that symbols *σ_i_* ∈ Σ were hypothesized to be recruited independently from the same spatial afferents as spatial symbols *δ_k_* ∈ Δ. In turn, this means that they might not be temporally co-active with *γ*^(*s*)^. Therefore, the temporal integration window for dom_Σ_ of *γ*^(*s*)^ needs to increase. Moreover, the symbols need to be presented in temporal order to preserve their spatio-temporal relationship. That is, suppose there exists a path of *N* symbols *σ*_1_ *, …, σ_N_*. This means that symbols *σ_i_* and *σ_i_*_+1_ are not only spatially related, but also temporally adjacent.

Activating symbols *σ_i_* ∈ Σ of a path in their temporal order induces a linked list in time, depicted in Figure 6A. Then, increasing the temporal integration window from one scale to the next follows immediately according to Subsection 3.2 and Subsection 3.3.

**Figure 6:**
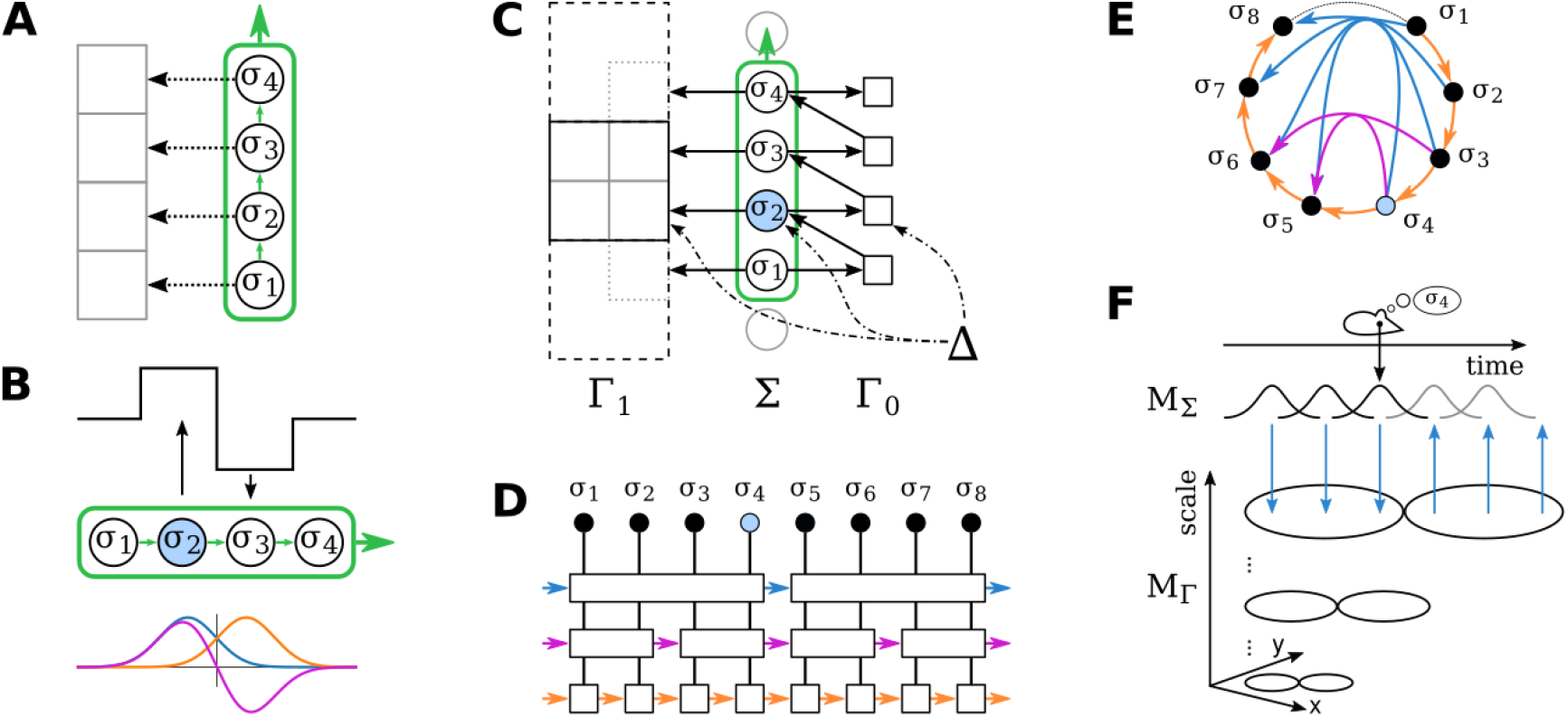
On-line learning of the Transition Scale-Space with temporal buffering. (A) Consecutive activation of symbols *σ_i_* ∈ Σ induces a linked list in time (green box). This preserves temporal (small green arrows) and spatial structure (indicated as arrows to discrete spatial areas). (B) Receptive fields of transitions for temporal integration as Haar wavelet (discrete case, top row) and Difference of mean-shifted Gaussians (biologically plausible representations, bottom row) on scale 0. Temporal transition start and end indicated as black arrows. (C) A transition on scale *s >* 1 (left) needs to associate its on-center (black solid box) with multiple symbols of Σ (middle). To keep their spatio-temporal structure, they are activated in sequence (green box) by iterating scale *s* = 0 of the scale-space (right). Black dashed box indicates spatial off-center (partial area only for visual purposes). (D) Simplified depiction of (C) for three scales. (E) The temporal buffer can be implemented as circular buffer. (F) Multi-scale illustration for biologically plausible receptive afferents. For further explanations for each panel, see Subsection 3.5.

Specifically, for the discrete case, the receptive field for the temporal integration can be modelled as a Haar wavelet, illustrated in the top row of Figure 6B. The positive part of the wavelet corresponds to the start of the transition between symbols of Σ, and the negative part its end. To remain consistent with the size increment of the spatial receptive field from one scale to the next, the temporal integration window thus has to increase by a factor 2.

Now consider biologically plausible representations. Likewise spatial afferents in Subsection 3.3, the activity of a symbol *σ* ∈ Σ is assumed to be Gaussian. Then, the temporal receptive field of a transition *γ* ∈ Γ can also be modelled as Gaussians, for instance as the first derivative of a Gaussian, or as a difference of mean-shifted Gaussians (see bottom row of Figure 6B for an illustration of the latter). The scaling of the temporal receptive field can thus be derived analytically according to Equation 4. Conclusively, the variance of the temporal integration window doubles, which means, following the same arguments as in Subsection 3.3, that its width increases according to 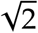 from one scale to the next.

It remains to understand how symbols *σ_i_* ∈ Σ can be presented to transitions *γ*^(*s*)^ ∈ Γ on scale *s >* 0 in temporal order and in a suitable temporal window. It is unlikely that spatial inputs activate symbols *σ_i_* in the correct manner. The reason is that symbols *σ_i_* ∈ Σ were hypothesized to be recruited on spatial afferents Δ and that only one (or a few) symbols *σ_i_* ∈ Σ are active at any single place (Waniek, 2018). That is, spatial input will activate only singular symbols. However, observe that scale *s* = 0 of the TSS learns transitions on singular elements of both Δ and Σ. Then, iterating the smallest scale of the TSS will induce a suitable temporal buffer.

More precisely, suppose that spatial inputs from Δ arrive at a given frequency at Σ and Γ. This frequency determines a time window *T*_Δ_ between consecutive inputs from Δ. Further assume that *T*_Δ_ is longer than the time it takes to iterate the transition system on the smallest scale. Then, it is possible to nest multiple such iterations in *T*_Δ_. This creates a spatio-temporally ordered list of symbols *σ_i_* ∈ Σ, or in other words, a temporal buffer. In turn, the temporal buffer can be used to associate higher scales. This process is depicted in Figure 6C with interactions between Σ and the two spatial transition scales Γ_0_ and Γ_1_. Figure 6D shows a simplified illustration for three scales. Changing the illustration such that the ends of the temporal buffer are connected reveals that this form of temporal buffering induces an oscillation around spatial input from Δ, depicted in Figure 6E. In other words, it can be implemented as a ring buffer that gets updated by novel inputs arriving from Δ. Figure 6F illustrates the interaction between temporally adjacent symbols and the spatial scale-space.

### 3.6 Sequence planning and shortcut discovery with Transition Scale-Spaces

This section answers how an animal can produce sequences between two arbitrary symbols *σ_s_* and *σ_t_* using a scale-space representation *ℳ*^S^. The presented multi-resolution approaches are similar to a method that was previously described by Behnke, 2004 in the domain of robotic navigation, and related to the hierarchical look-ahead model presented in Erdem and Hasselmo, 2014; Erdem, Milford, et al., 2015. Moreover, the described methods are graph search algorithms and based on well-known techniques such as Dijkstra’s algorithm (W. Dijkstra, 1959) or flooding algorithms (Fishkin and Barsky, 1985; Lynch, 1996; Silvela and Portillo, 2001).

As in the previous sections, transitions are treated irrespective of the transition bundle to which they belong. This simplifies the description without violation of the logical considerations of MTS and, thus, improves clarity of the presentation. Likewise, the temporal transition memory *M*_Π_ will be omitted. Finally, assume that the memories already formed, i.e., the scale-space was already built and corresponding symbols assigned to transition images and domains on each scale.

Consider an *ℳ*^S^ with only a single scale. Finding a sequence from a symbol *σ_s_* to *σ_t_* requires only few operations. First, *σ_s_* must be activated in *M*_Σ_. Then, a corresponding transition *γ_i_* that is defined for *σ_s_* will be activated in *M*_Γ_. In turn, *γ_i_* activates a set {*σ_j_* } of subsequent symbols *σ_j_* to which *γ_i_* leads to. The combination of these two steps is called *expansion*. If there exists at least one path from *σ_s_* to *σ_t_*, recursively repeating this process will eventually activate *σ_t_* in *M*_Σ_. Obviously, this is a flooding algorithm applied on a graph (Lynch, 1996; Silvela and Portillo, 2001), where nodes correspond to symbols *σ_i_*, and transitions are made explicit by transition encoders *γ_j_*

During the expansion step, any expanded symbol needs to remember its parent symbols. For instance, if symbol *σ*_6_ activated transition encoder *γ*_0_, which in turn activated *σ*_7_, then *σ*_7_ needs to remember *σ*_6_ as its parent symbol. Then, after successful activation of the target symbol *σ_t_*, a viable sequence from *σ_s_* to *σ_t_* can be found by backtracking from *σ_t_* via any of the parent symbols. This is the backtracking step of Dijkstra’s algorithm, modified to select an arbitrary parent from the set of feasible parents. Note that by following the symbolic sequence *σ_s_* ↝ *σ_t_*, the spatial sequence can be inferred because transition encoders *γ_i_* learned transitions in the symbolic space Σ and in the spatial input space Δ jointly.

The animal can use multiple scales in several ways. One possibility is to apply the expansion and backtracking algorithms only on a behaviorally necessary level. For instance, if the animal needs to find an exact sequence, then it should use the lowest level of *M*_Γ_ because it contains the highest resolution. If, however, the animal needs only an approximate direction towards a goal or when computational time is limited, it can operate on the highest available level. Intuitively speaking, the highest level represents approximate knowledge and, thus, can be used to find coarse estimates. Another possibility is to use all scales during sequence planning, which can be implemented in different ways.

One variant is *descending mode*, in which the sequence is first approximated on the highest available scale. After finding the target symbol *σ_t_* on the highest scale, backtracking to the starting symbol *σ_s_* yields sub-goals. These sub-goals are defined as all those symbols that are reachable via only one transition on this scale. Then, the sub-goals are used on the next smaller scale to refine the trajectory, illustrated in Figure 7. In both panels of the figure, the start symbol is *σ*_3_ (blue color) and the target is *σ*_6_ (green color). Panel A shows two scales in a one dimensional example and includes all details about the transition encoders 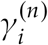, where *n* is the scale. That is, a transition encoder 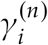 (black box) learns transitions simultaneously in the symbolic space Σ as well as the spatial input space Γ, indicated by white boxes containing corresponding symbols. The expansion and backtracking steps on scale 1 yield intermediate symbols, shown demonstratively only for *σ*_0_ (pink color). Then, the sequence is refined via scale 0 that computes only the trajectory from *σ*_3_ to the sub-goal *σ*_0_. Note that during retrieval, transition encoders 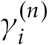 don’t need access to spatial symbols *δ _j_*. However, these spatial symbol can be inferred due to activation of the transition encoders. Panel B introduces a reduced graphical depiction of this process, in which details about transition encoders are omitted. The panel also illustrates that the integration areas of higher scales don’t need to perfectly align at the borders with smaller scales. Figure 7C-E present an illustration of this process for a two-dimensional example in a grid-world. An animal that wants to compute a trajectory from its current location to a target (cheese) first retrieves a coarse estimate that activates multiple symbols in a coarse target region (panel D, purple small squares) and intermediate sub-goals (pink stars). Backtracking effectively reduces the search space for the refinement process. That is, symbols that are not on the coarse path (panel D, gray circles) don’t need to be expanded on the next finer scale (panel E).

**Figure 7:**
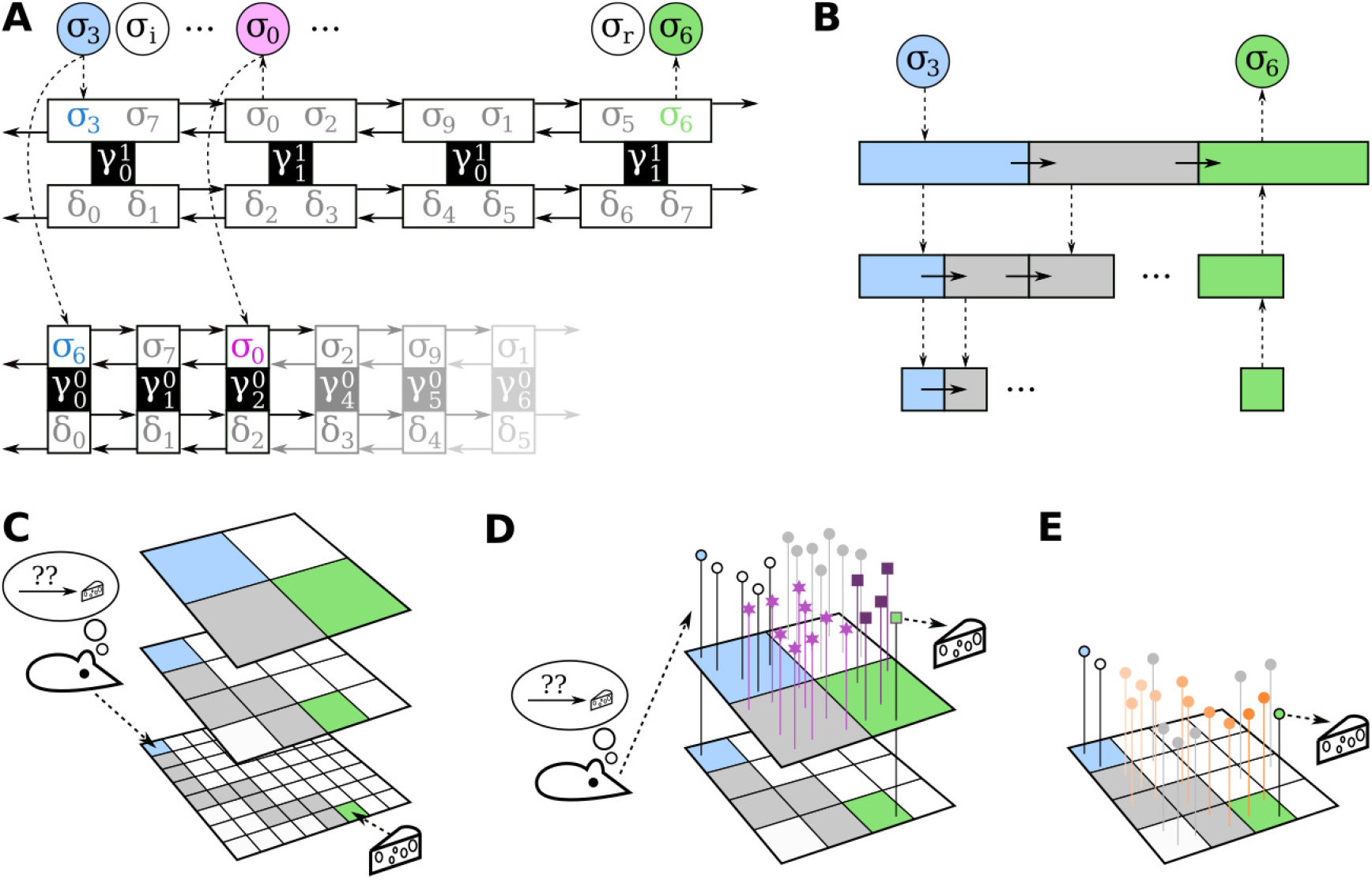
Retrieval of sequences with a scale-space in *descending mode*. (A) 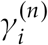 learn transitions in two spaces simultaneously (cf. Figure 3). During retrieval, one space is sufficient (dashed lines) and expansion starts on the highest level from start (blue circle) to goal (green circle). After an approximate solution was found on a level, sub-goals are generated (magenta circle). Then, the next lower scale computes expansions to these sub-goals. (B) Simplified depiction of (A) that omits details about the input spaces of transitions. (C) An animal wants to compute a path from its current location to a target using a Transition Scale-Space. (D) It can first compute an estimate on the highest scale, prune unnecessary symbols (gray circles), and generate sub-goals (magenta stars). (E) The trajectory is subsequently refined by planning on a finer scale to these sub-goals. Note that, in the discrete case, diagonal movements violate constant cost assumptions of MTS (see Waniek, 2018). In case of multiple viable trajectories, the algorithm chooses one arbitrarily.

Another variant is *ascending mode*, in which the search for *σ_t_* starts on the lowest scale. After a number of failed attempts to discover *σ_t_* on this level, the search continues on a larger scale. This is repeated until the target is found, which may eventually, but not necessarily, involve the highest scale. In other words, this automatically adapts the scale to the real distance to the target. In contrast to *descending mode*, this variant requires the backtracking operation only once the target is found.

In both modes, the animal has access to the first steps of a detailed sequence after only a brief computational time, and can refine this sequence while already moving. Note that there might be multiple feasible ways to expand a path on each scale in either mode. In this case, the algorithms used in Section 4 and presented in Appendix C randomly select one possible solution during backtracking. Finally, both modes have the same amortized runtime complexity. Specifically, assume a planning operation with a TSS to a remote location, and that this location is significantly further away than size of domains of the largest scale. Then, the runtime will be dominated by the number of retrievals on the highest level of the TSS in either retrieval mode, and the cost of expansions on lower scales are negligible. Pseudo-code and further details of the algorithms and sub-routines can be found in Appendix C.

Conclusively, the runtime issue that was motivated in Subsection 3.1 can be solved by an appropriate number of levels in a TSS.

## 4 Experiments

The scale-space data structure and algorithms were examined using simulations of a virtual agent on a plane surface. In particular, the *ℳ*^S^ that was simulated consists of a memory *M*_Σ_ that stores symbols *σ_k_*, *k* ∈ *N*_Σ_ and a memory *M*_Γ_ of spatial transitions.

If not stated otherwise, symbols and domains and images of transitions were computed in an offline manner. That is, transition centers were distributed hexagonally at the beginning of a simulation because of results presented in Waniek, 2018. Then, symbols were distributed randomly in the environment. Subsequently, images and domains of transitions were computed from the relative locations between symbols and transitions. Specifically, each transition pooled all symbols according to a nearest neighbor search, and lead to all symbols in its image simultaneously (see Figure 2 for reference). Thereby, transitions discretize the input space and perform Voronoi clustering.

The source code to reproduce all data for the figures and results presented in this section are available online under an Open Source License^1^.

### 4.1 Worst case timings on a linear track

Segment trees or interval skip lists are widely used and their characteristics well known. The objective of this experiment was thus to simply show the acceleration capabilities of the data structure to find an element in the worst-case scenario.

A simulated agent was put on a linear track of length 10 m to compute the number of recursive retrievals that are required to find a specific element. Spatial symbols were distributed on the track quasi-randomly using the Hammersley point set, a low-discrepancy sequence. Then, a Transition Scale-Space (TSS) consisting of 7 scales with periods 0.2, 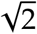 · 0.2, 0.4, 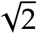 · 0.4, 0.8, 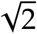 · 0.8 and 1.6 m between transition encoders was computed offline by associating symbols to transition encoders using a nearest-neighbor search. The starting symbol closest to the start of the track was designated as *σ_s_*, and the symbol closest to the end as *σ_t_*. The numbers of necessary recursive retrievals from *σ_s_* until *σ_t_* was found are presented for each scale numerically in Table 2.

**Table 2:** Number of required recursive expansions until a target symbol *σ_t_* is found on a linear track.

|  |  |  |  |  |  |  |  |
| --- | --- | --- | --- | --- | --- | --- | --- |
| Grid Period | 0.2 | $\sqrt{2} \cdot 0.2$ | 0.4 | $\sqrt{2} \cdot 0.4$ | 0.8 | $\sqrt{2} \cdot 0.8$ | 1.6 |
| Recursions | 50 | 35 | 25 | 18 | 12 | 9 | 6 |

### 4.2 Wave propagation dynamics in two dimensions

In two dimensional settings, the expansion procedure is expected to resemble a propagating wave of activity when no additional restrictions on the expansion are imposed. To examine and illustrate this behavior, expansion was computed according to Algorithm 2 in a square environment of size 1 × 1 m. A total of 400 spatial symbols were distributed either randomly with a minimal distance of 0.02 m or according to the Hammersley point set. The objective of the latter was to observe any difference in behavior with more regularly placed symbols. Then, transitions were computed for a period of 0.2 m between transition encoders offline. The expansion was started at symbol *σ_s_*, which was defined to be the closest symbol to position (0.2, 0.2), and stopped at the target symbol *σ_t_*, closest to position (0.8, 0.8). After expanding and localizing *σ_t_*, 300 Monte Carlo samples of symbolic sequences were computed using backtracking and randomly selecting a parent during each backtracking step.

Figure 8 shows the results of the expansion and the emerging propagating wave from *σ_s_* to *σ_t_*. Black dots indicate symbols which are currently active during the expansion. The final expansion panel of each row shows a solution space as gray hexagons and Monte Carlo samples as blue lines. The solution space is given by the sequence of spatial domains of subsequently activated transitions on a solution sequence and forms a channel from start to target.

**Figure 8:**
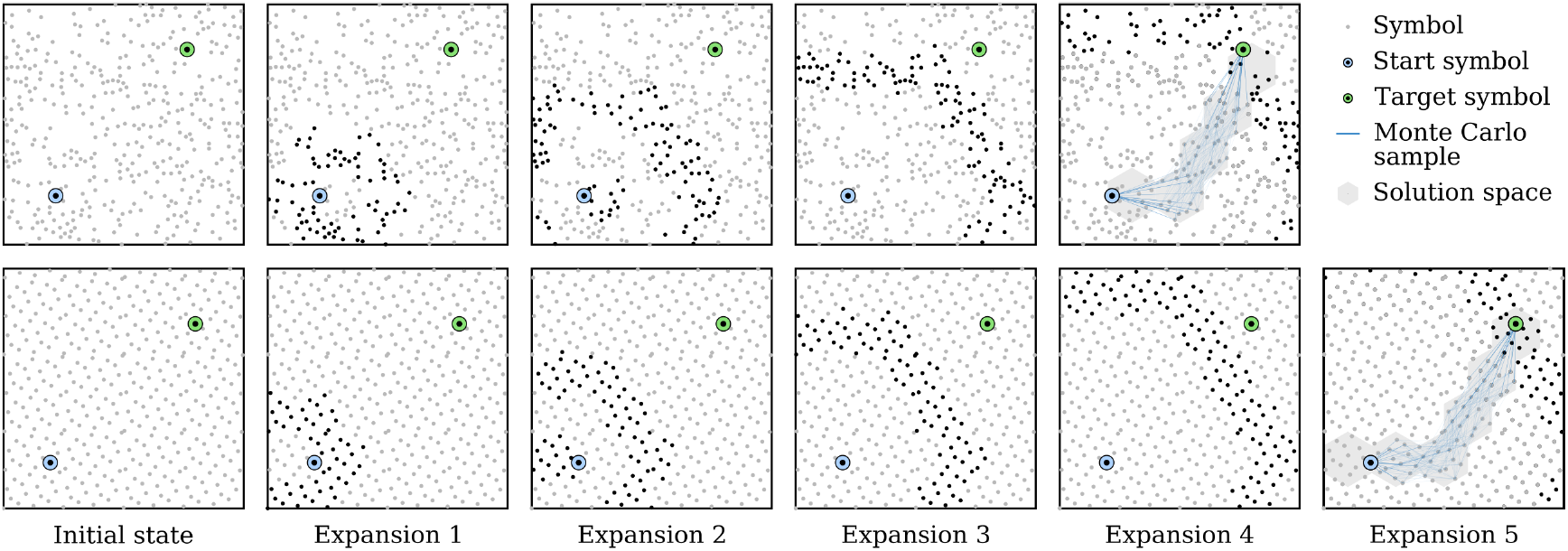
Propagating wave during expansion from start to target in a square environment. Top row shows results for randomly distributed symbols (gray dots), bottom row for symbol coordinates according to the Hammersley point set. Transition encoders were placed regularly on a hexagonal lattice (not shown) due to results presented in Waniek, 2018. Final panels include the solution space (gray area) and 300 Monte Carlo samples (blue lines). The solution space are all those images of transition encoders which contain symbols that are on a shortest path from goal to target. See Appendix E for results in a non-Euclidean space.

The only distinctive qualitative difference between randomly and more regularly placed symbols is the form of the propagating wave. When the symbols are regularly placed, then the wavefront is hexadirectional during later expansion steps due to the perfectly regular placement of transition centers. For randomly placed symbols, the wavefront is more tattered, but still resembles the hexadirectional characteristic of the regular case. The difference in the solution space and the number of expansions is due to the selection of the start and target symbols and, thus, negligible.

Note that more than just one solution space may exist due to the underlying assumption of MTS. Specifically, MTS assume a constant-cost operation to move from the domain of a transition to its image. However, only one solution space is illustrated for each example in Figure 8. Also note the active symbols close to the start symbol *σ_s_* during the second expansion. This behavior is because these symbols were not co-activated during start even though they are in close vicinity to *σ_s_*, and because the implemented transition system does not include any knowledge about spatial directivity of the expansion.

### 4.3 Scale-space planning with sub-goal generation in descending mode

The descending mode of trajectory refinement was analyzed using a rectangular area of 2 × 5 m^2^. In this area, 250 symbols were distributed randomly, keeping a minimal distance of 0.05 m between each pair of symbols A TSS consisting of 5 scales with periods 0.2, 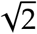 · 0.2, 0.4, 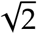 · 0.4 and 0.8 m between transition encoders was then computed offline.

The symbol closest to coordinate (0.4, 0.1) was selected as start symbol *σ_s_*, and the symbol closest to coordinate (1.8, 4.8) as target symbol *σ_t_*. Qualitative results are illustrated in Figure 9. The search starts on the highest available scale, which computes an entire approximate solution from start to target. The solution space of this scale is indicated by gray hexagons in the leftmost panel of the figure. Also, this generates sub-goals, indicated by magenta circles.

**Figure 9:**
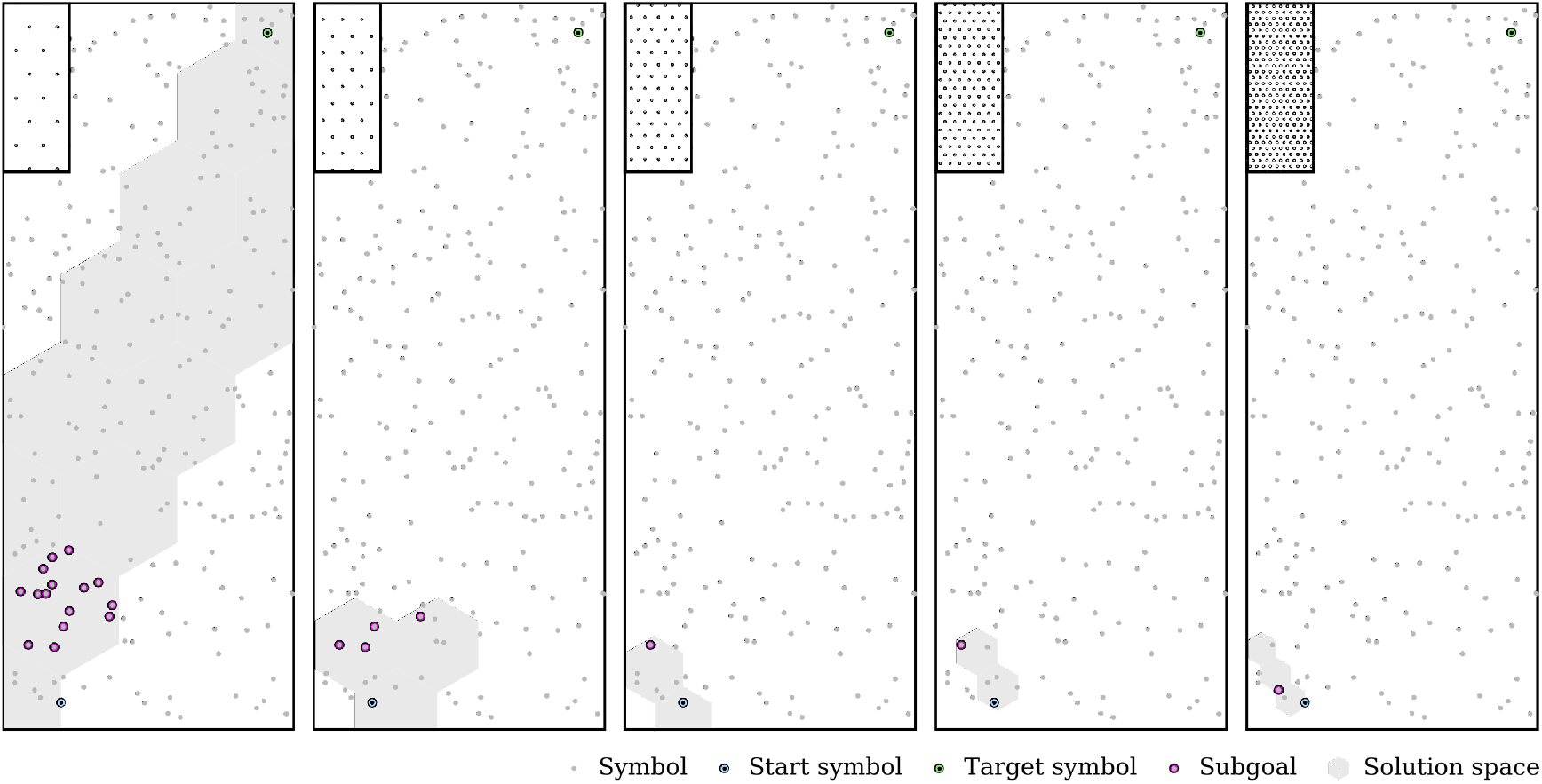
Searching a viable sequence from start (lower left) to target (upper right) using descending mode in a space with randomly sampled symbols (gray dots). The search from start (blue symbol) to target (green symbol) starts on the highest available scale (left panel) and generates sub-goals (magenta symbols). Gray area displays the solution space, which consists of all images of spatial symbols that were expanded and contain trajectories with equal distance. The solution is then refined using finer scales, which compute only partial solutions to the sub-goals from the next higher scale (remaining panels). Distributions of transition encoders are shown in the small inlays in each panel, and were precomputed based on results presented in Waniek, 2018.

After defining sub-goals, the procedure drops down to the next scale and computes trajectories only to the previously determined sub-goals. This limits the search space on smaller scales. The process repeats until the smallest scale is found, which is illustrated in the other panels of the figure.

### 4.4 Scale-space planning in ascending mode

The qualitative behavior of the ascending mode was examined likewise the descending variant. That is, 250 symbols were placed randomly in an environment of 2 × 5 m^2^, with a minimal distance of 0.05 m between symbols. Then, a TSS consisting of 5 scales with periods 0.2, 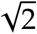 · 0.2, 0.4, 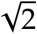 · 0.4 and 0.8 m was computed offline.

The initial symbol was determined by finding the closest symbol to coordinate 0.4, 0.1, and the target by the symbol closest to coordinate 1.8, 4.8. During search, 3 expansions were computed on each scale before ascending the search to the next scale.

An example of the ascending mode is depicted in Figure 10. Each column shows the expansion on one scale. The first scale searches locally in the area that is indicated by the gray hexagons. Subsequent scales expand the respective search radius, and propagate the search outward until the target is found on the highest scale. For each scale, black dots indicate the symbols that were active at the end of the search on this scale. After finding the target symbol, 100 Monte Carlo trajectory samples were drawn by backtracking from the target symbol to the start symbol, and selecting a random parent during each backtracking step. Note that this results in small trajectory segments and low variability close to the start of the trajectory, but larger segments and higher variability close to the target. In the figure, these samples are illustrated as blue lines on top of the search of the largest scale.

**Figure 10:**
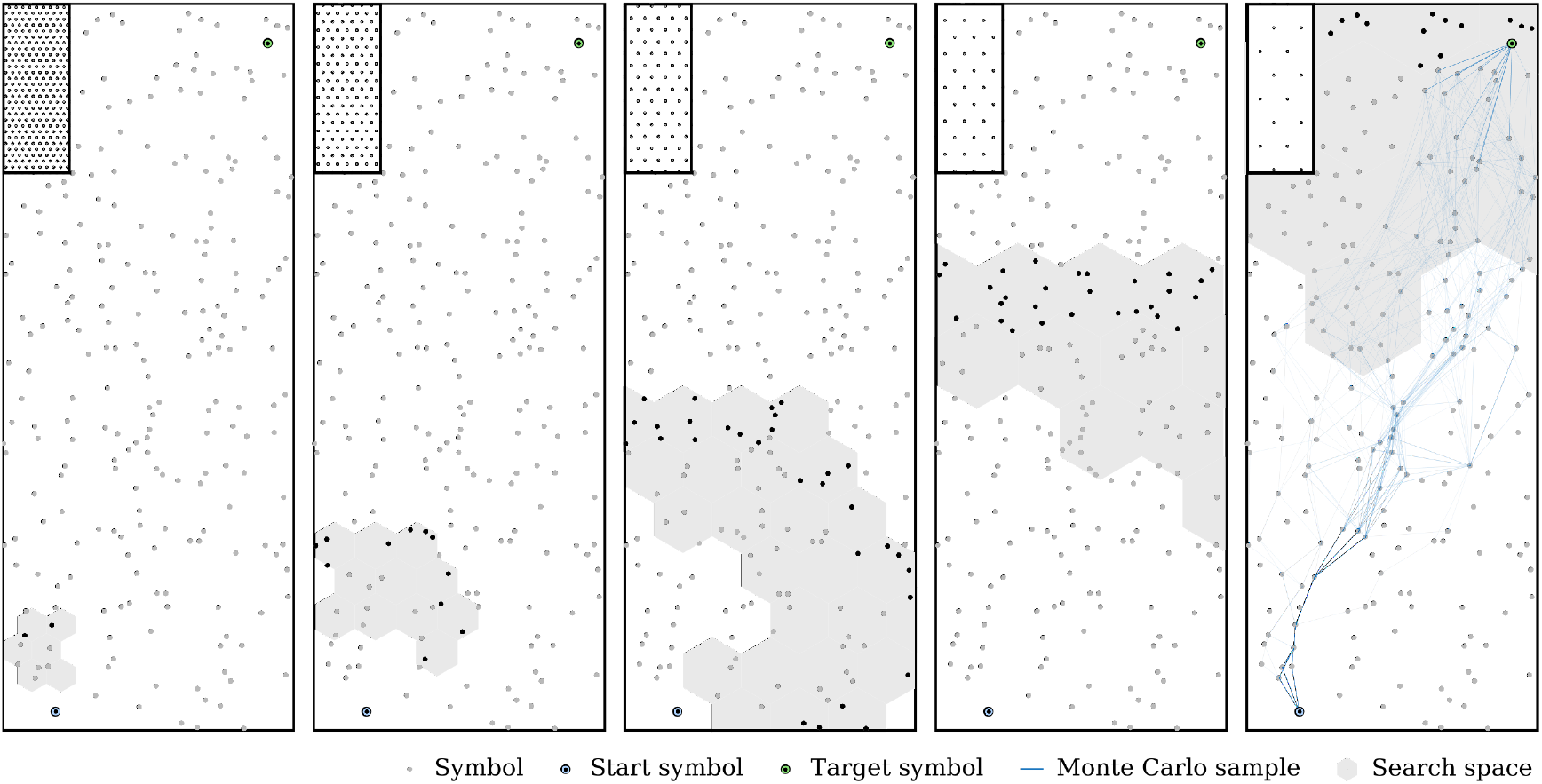
Searching a viable sequence from start (lower left) to target (upper right) using ascending mode. Each scale is expanded at most 3 times before ascending to the next scale (panels from left to right), until the highest scale is reached. Note that, in contrast to Figure 9, here the gray area depicts all images of spatial transitions that were expanded during search, and not the solution space. Black dots indicate the last symbols that were active on a scale before ascending to the next scale or when the target was encountered. The rightmost panel includes Monte Carlo samples (blue lines), generated using the backtracking algorithm. Distribution of transition encoders is shown in the small inlays in each panel, and was precomputed according to prior theoretical results presented in Waniek, 2018. Symbols (gray dots) were distributed randomly on the input space.

### 4.5 Shortcut discovery in the Morris water maze experiment

A simulated version of the Morris water maze experiment was used to examine the capability to find shortcuts in spatial information using a TSS. Qualitative results are presented in Figure 11 for three independent simulations. In each simulation, an agent started from the initial location (blue square) and traveled the environment with a radius of 2 m until the target platform (green circle) was found using movement statistics similar to real rodents. Symbols (or place cells; gray dots) were distributed along the recorded trajectory (black line) as follows, inspired by Growing Neural Gas (GNG)(Fritzke, 1995). As soon as the agent was further away than *m*_dist_ = 0.05 m from the closest symbol, a new symbol was recruited. To introduce stochastic symbol placement, this novel symbol was associated with a coordinate that was drawn from a unit circle centered at the current position of the agent and normally distributed with a standard deviation of 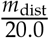, where the denominator was chosen arbitrarily. Subsequently, a TSS with 5 scales was computed offline for grid periods of 0.2, 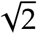 · 0.2, 0.4, 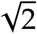 · 0.4 and 0.8 m (distribution of centers shown in the legend of the figure). Specifically, each symbol was associated with the domain of the closest transition encoder, and to the image of the surrounding encoders.

**Figure 11:**
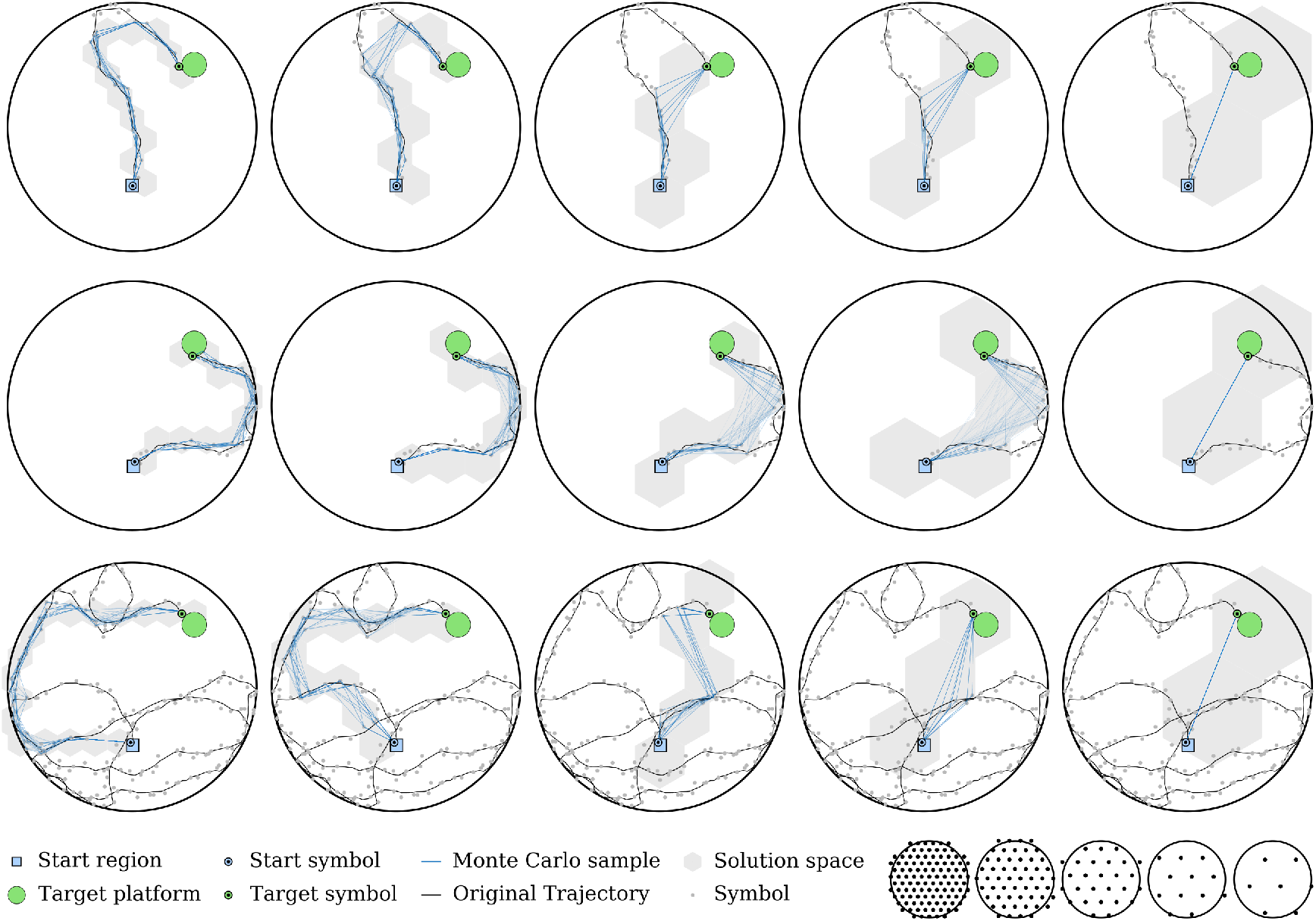
Results for three simulated Morris water maze experiment. Initially, the agent performed a random walk from Start (blue square), until it found the target platform (green circle). Meanwhile, novel symbols *σ_i_* ∈ Σ (gray small points) were spawned based on distance to previous symbols. The original random trajectory that the agent followed is shown in all plots as black line. Subsequently, possible solutions were computed for each scale individually (one scale per column). Each row represents one experiment, each column using one specific scale to compute the solution space (gray hexagons) and 100 candidate solutions (blue lines). The legend shows the distributions of transition encoders for each scale, which were precomputed according to prior results and assumed constant over the course of the simulation. See Subsection 4.5 for further details.

For each scale individually, the solution space was computed according to Algorithm 2. That is, starting from the initial symbol, a sequence of viable transitions and symbols was computed until activation of the target symbol, which was defined as the symbol closest to the target platform. Figure 11 shows the solution space as gray hexagons. For each scale, 100 Monte Carlo samples (blue lines) were computed by backtracking from the target symbol to the initial symbol and selecting a parent symbol randomly during each backtracking step.

The figure shows that, on the smallest scale, reproduced sequences and the solution space remain close to a shortest path on the original trajectory. On higher scales, sample sequences and the solution space exhibit shortcut characteristic, essentially crossing parts of the environment that were not explored previously by the agent.

## 5 Discussion

Motivated by the runtime complexity of sequence retrievals in transition systems, this paper introduced the Transition Scale-Space (TSS). The TSS is an abstract data structure that contains knowledge about transitions between symbols on multiple scales, shares several properties with the rodent Hippocampal formation, and in particular MEC. Most important, Subsection 3.3 and Subsection 3.5 showed that, given the assumption that they follow a Gaussian tuning curve, the optimal scale increment for both spatial and temporal receptive fields between consecutive scales of a TSS is 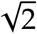. Consequently, it is hypothesized that the entorhinal-hippocampal loop forms a TSS, and that individual grid fields are spatio-temporal kernels that learn transitions both in spatial inputs, as well as a symbolic space. While the first is presumably given by spatially modulated input such as boundary-vector-cells, head direction cells, or speed cells, the latter is represented by place cells. Moreover, bottom-up learning of a TSS needs a mechanism to temporally buffer the symbolic input space. In total, this yields the TSS model for grid cells, depicted in Figure 12.

**Figure 12:**
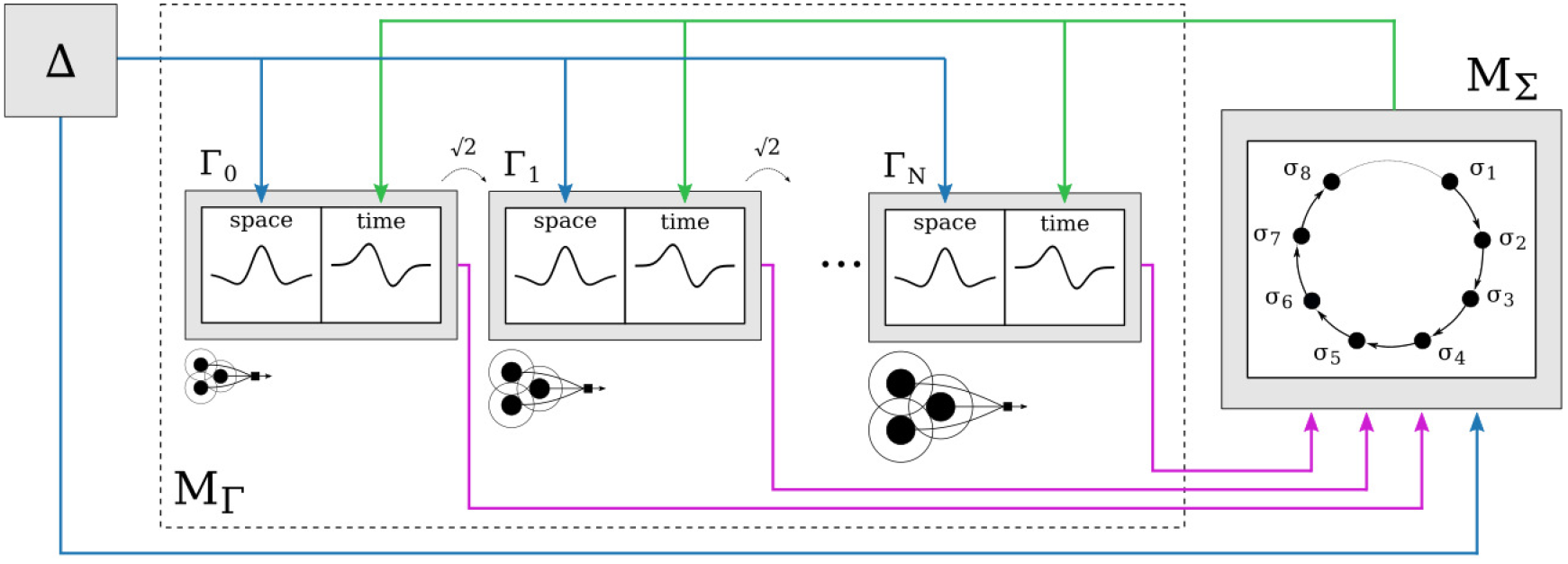
Transition scale-space model for grid cells with memories *M*_Γ_ (dashed box, middle), *M*_Σ_ (gray box, right), and spatial afferents Δ (gray box, left). It is hypothesized that each grid field of a grid cell is a space-time-kernel (gray small boxes within *M*_Γ_) that represents spatial transitions. The spatial part of the receptive field learns transitions on input space Δ (blue arrows), which is presumably given by boundary-vector-, head direction, speed cells and other spatially modulated inputs. The temporal part integrates temporally ordered activity in a symbolic space Σ, itself hypothesized to be represented by place cell activity. For accelerated searches and retrievals of sequences, *M*_Γ_ consists of *N* different scales Γ*_i_* (multiple scales depicted as horizontally arranged small gray boxes). For biologically plausible representations, the space-time kernels increase optimally by 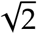 from one scale to the next (dashed arrow between small gray boxes). Learning multiple scales requires temporally buffered access to symbols *σ_i_* ∈ Σ (ring buffer within *M*_Σ_), which allows to relate the model to Theta phase precession (see Subsection 5.2).

Although the results were presented in the context of spatial navigation due to the involvement of grid and place cells in navigational tasks, the results apply to arbitrary transition systems that operate on topological data. Now, these results will be discussed concerning their plausibility, predictions, and relation to other work.

This section is organized as follows. First, the data structure, algorithms, and results of the previous sections will be critically discussed with respect to biological plausibility, and testable predictions derived from TSS. Subsequently, temporal buffering and its potential relationship to Theta and TPP will be examined. This is followed by a detailed discussions about similarities and differences to other models for grid cells. Finally, TSS will be related to other areas of research, in particular to results from signal processing, mobile robotics, and computer science.

Note that, throughout the section, the terms symbol and neuron may be used interchangeably. Moreover, and if not stated otherwise, the term TSS will be used inclusively to mean both Multi-Transition System (MTS) and the extension to multiple scales presented in this paper.

### 5.1 Scale-space computations and biological plausibility

The properties and dynamics of TSS are compatible with observations from the Hippocampal formation. For instance, the activity of symbols in a TSS forms a traveling wave during expansion from start to target, depicted in Figure 8. Also, the Hippocampal formation shows traveling waves of activity (Lubenov and Siapas, 2009; Patel et al., 2012). However, TSS was not presented as a spiking neural network but in mathematical and algorithmic form. Thus, aspects of TSS will now be examined with respect to their biological plausibility. This will also be used to derive testable predictions from the mathematical model.

Figure 8 shows that, during expansion, TSS activates all intermediate symbols in form of a traveling wave until the target is reached. This is unlikely to occur in a biological neural network, as this could lead to saturation of activity in the network. A solution to this shortcoming of the model is to activate only a sparse subset of symbols in the image of each transition during path expansion. However, this then involves the danger to miss the target symbol. To avert the latter problem, the target should be overrepresented by several symbols. Essentially, this increases the chance to activate one of those symbols that corresponds to the target. It is expected that the number of symbols that are used for such an over-representation is inversely proportional to the probability of a symbol to activate during transition evaluation and to the importance of the goal. However, future work is required to examine if this conjecture holds or not. Note that the idea to over-represent targets is consistent with evidence that – in rodents – place cells accumulate near goal locations (Boccara et al., 2019; Dupret et al., 2010; Hollup et al., 2001; Lee et al., 2006; Mamad et al., 2017).

Table 2 shows the decrease in computational time to find a certain symbol. Moreover, Figure 9 and Figure 10 show that a TSS can be used to quickly compute approximate sequences. Importantly, the first steps of the retrieved sequences are *valid* according to the definition given in Waniek, 2018, which means that they are consecutive and connected and thus allow deterministic movement. Valid sequences might be of importance during tasks that require precise movement along previously acquired places. On the other hand, approximate solutions could also be used directly. This appears to be of utmost interest in other behavioral tasks. For instance, consider the Morris water maze experiment, in which a rodent has to establish a route to reach the target platform. Waiting for an accurate computation to finish is not an option for a rodent, given the perilous situation in which it is during this experiment. Rather, the animal requires an algorithm that quickly yields an estimate of the solution so that it can start moving, and refine the trajectory online. The qualitative results shown in Figure 11 demonstrate that a TSS provides such an algorithm. Using a large scale quickly narrows the search space, or is even able to present a direction directly towards the goal. Note that a large scale comes at the cost of an increased variance of the solution, indicated by the large gray areas in the figures. Conclusively, it is expected that rodents operate not only on the highest available scale, but either in ascending mode to retrieve sequences that are initially valid, or on all scales in parallel.

The TSS and the simulation results were not presented in form of spiking neural networks. However, a neural network implementation of a TSS is conjectured to require only a comparatively simple neural micro-circuit. Recall Subsection 3.2 and Subsection 3.3 which explain that each spatial transition of a TSS represents transitions as intervals. Moreover, each level of the hierarchy is conceptually identical to other levels and differs only by the size of the interval. Then, learning that a symbol falls into the domain of a spatial transition on a higher scale follows exactly the same rules as lower scales and, thus, generalizes well ^2^. Moreover, representing associativity between transitions and symbols requires only simple binary logic elements, as discussed previously in Waniek, 2018. Thus, a neural network implementation of a TSS needs to only differ in the size of the spatial and temporal sampling process on each scale. Finally, the expansion algorithms that were used during the simulations can be parallelized for a distributed message passed system, and thus appear to be suited for spiking neural networks.

Both expansion methods, descending and ascending, as described in Subsection 3.6, lead to two different waves of activity. The first, already discussed above and presented in Figure 8, is a wave of activity from start to target. The second is a wave of activity that propagates along the hierarchy of TSS. Figure 9 shows that, during descending mode, the second wave starts on the highest (or coarsest) level of TSS and, after determining sub-goals, activity moves successively to lower scales. In contrast, the second wave starts on the lowest level and moves to higher scales, as depicted in Figure 10. In particular the latter is compatible with evidence from the Hippocampal formation. Not only increase grid fields systematically along the dorsal to ventral axis (Brun et al., 2008; Stensola et al., 2012), also activity propagates in the form of a traveling wave along the axis (Lubenov and Siapas, 2009; Patel et al., 2012). This is also compatible with the observation that increased temporal integration windows of transitions on higher scales necessarily lead to temporal lag, as will be discussed further below.

Both modes were used to quickly retrieve approximate solutions with detail only for the next steps. The modes correspond to usage of multi-resolution data structures that found widespread adoption in other areas of computer science (Behnke, 2004; Finkel and Bentley, 1974; Kambhampati and Davis, 1986; Lu et al., 2011; Orenstein, 1982). In particular the computer graphics community characterized the performance benefit that comes with multi-resolution representations and relies heavily on acceleration data structures of this form (MacDonald and Booth, 1990; Wald et al., 2009). Intuitively, large scales of such representations can be used to quickly prune data and select only a subset of the original data. Successive use of finer scales then refines the data to yield only singular elements of interest. This also applies to TSS in the sense that higher scales help to quickly retrieve feasible trajectories, and leads to the following prediction. Disruption of all larger scales of a TSS will lead to decreased performance during retrieval. Specifically, if there is just a single scale, then the TSS will suffer from the computational times that motivated the development of TSS, presented in Subsection 3.1. This could be shown in an experiment in which animals have to navigate to very remote targets. Targeted lesion or inhibition of grid cells with large grid fields should lead to increased computational time during planning, measurable as an increase of the temporal delay between onset of pre-play and movement. Moreover, the number of failed navigation attempts should increase. The latter is because the activity of the traveling wave from start to target might diffuse due to noise of neural activity. In addition, increased navigation failure is also likely due to reward diffusion during sequence selection, an issue that was addressed previously in the model by Erdem and Hasselmo, 2014.

Sequence selection was computed using classical backtracking in the results presented in Section 4. In this form, the presented algorithm is biologically implausible as it would require reverse information propagation. However, there are likely candidates for alternative implementations. First, recall that the data within a TSS represents a connected graph with constant edge weights, and searching a trajectory corresponds to finding a shortest path in a weighted graph. In turn, this edge weight can be interpreted as a temporal delay for spike propagation between vertices of the graph. Then, computing a winning trajectory could be achieved by reducing the temporal delay between vertices that activate the goal location. Iterating this is expected to successively compress expansion of a path *in time*. The outlined process describes the computation of an eligibility trace, known from RL, and is also related to reward diffusion that was used in Erdem and Hasselmo, 2012 and Erdem and Hasselmo, 2014. Also, this mechanism could explain why rats appear to pre-play multiple similar trajectories during homing tasks, observed by Pfeiffer and Foster, 2013. More specifically, repeated pre-play of similar sequences is expected to reduce the interspike interval between consecutive neurons that are on a sequence from goal to target. Analyzing temporal compression during pre-play will require significant amounts of data, and in particular simultaneous recordings of a large number of place and grid cells from a single animal.

### 5.2 Spatial pooling, temporal buffering, and Theta Phase Precession

Subsection 3.5 deduced that online-learning of a TSS requires temporally buffered symbols for spatial pooling. While a general requirement for buffering in scale-spaces was previously noted in the context of modeling visual receptive fields (Lindeberg, 2013), Subsection 3.5 suggests that this can be achieved *bottom-up* using iterations of the lowest scale of a TSS that are nested within a slower input cycle, illustrated in Figure 6E, as follows. Consider to learn spatial transitions between symbols on a larger scale, which means that symbols activate in a smaller portion of space than the sizes of the domain or image of a transition (see Figure 6C). To associate the correct symbols with the domain and image, Subsection 3.5 suggests to iterate the lowest scale of a TSS. More precisely, a recursive loop between *M*_Γ_ and *M*_Σ_ that evaluates transitions on the lowest scale effectively expands a path around a certain symbol, depicted in Figure 6E. Due to directionality of transitions, these symbols activate in a spatially coherent pattern. That is, this re- and pre-plays symbols in order that corresponds to their relative location to the current symbol similar to how place cells activate during Theta Phase Precession (TPP) (Schmidt et al., 2009; Skaggs et al., 1996). Then, all those symbols which temporally activate before the current symbol as well as the current symbol can be considered to be part of the domain of a transition and associated with it, while all those that follow can be considered to fall into the image and learned as transition targets (see Figure 6D and F for illustrations). Several predictions follow immediately from this description of temporal buffering. First, the temporal integration window of Spike-Timing Dependent Plasticity (STDP) of grid cells on larger scales must match their spatial receptive field for association learning with place cells and spatial afferents that fall into their receptive fields. Following Subsection 3.5, the temporal window ideally scales by 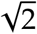 from one scale to the next. Second, targeted lesion of smaller grid scales should disturb subsequent larger scales. The reason is that such lesioning prevents the recursive loop between *M*_Γ_ and *M*_Σ_ on the affected scales. Consequently, the spatial coherence of the sequence of symbols is not guaranteed. Note, however, that temporal coherence between symbols is unaffected by such a lesion because, in an MTS, temporal transitions are learned in a separate memory *M*_Π_ (Waniek, 2018). Thus, to fully disrupt grid scales, it is expected that silencing of *M*_Π_ is also required. Third, there will be only a single scale in the absence of a temporal buffer or multi-scale representations of the input and symbol space. The reason is that if transitions of a larger scale project to symbols of smaller scales, or if symbols are only available in scales smaller than the transition scale, they need to be temporally available for learning with STDP as outlined above. If symbols are, however, available in larger scales, then transitions can be learned directly between symbols of corresponding higher scales. In either case, large scale transitions must not project to and activate smaller scale symbols during exploration, because this might activate all target symbols of the smaller scale simultaneously. In turn, this would disarray transition learning on that smaller scale and consequently lead to temporal and spatial incoherence.

As mentioned above, nested iterations of lower scales expand a path around a certain symbol, as depicted in Figure 6E. That means that starting nested iterations at a given symbol, for instance due to activation from Δ, will predict immediate next symbols. On the other hand, converging nested iterations from past symbols towards the given symbol repeats existing knowledge. To preserve linearity of spatio-temporal sequences, these nested iterations should have directional tuning and ideally only expand (or retract) symbols that are in the heading direction of the animal. Linearity of activation is well supported by data from rodent experiments (Jeewajee et al., 2013; O’Keefe and Recce, 1993). More precisely, experiments showed that, when an animal is running, place and grid cells that are along the animal’s heading direction perform TPP, which means that their firing activity progresses relative to Theta. It was previously noted that the phase reference that is provided by TPP is required to encode temporal order (Lisman, 2005). In addition, Theta modulation was previously used to recall episodic sequences of states in models of regions CA1 and Cornu Ammonis 3 (CA3) (Hasselmo, Bodelón, et al., 2002; Hasselmo and Eichenbaum, 2005). However, these models predate the discovery of grid cells and, thus, do not consider them. Note that conjunctive representations that have directional tuning, and which are supposed to be required for heading dependent linear temporal buffering, are not yet included in the TSS model. In addition, it is currently unclear how the inputs from the spatial input space Δ are optimally aligned with a temporal buffer.

TPP suggests that the alignment is ideally centered around the symbol that corresponds to the animal’s current location during constant running speed. Note that during TPP, cells in the heading direction of an animal spike in order of their temporal succession (Jeewajee et al., 2013; Schmidt et al., 2009). Setting this observation in context of TSS, it is hypothesized that Theta and TPP in particular are the observable effects of temporal buffering to learn and retrieve data from multiple scales. Moreover, it is conjectured that, assuming constant running speed, the optimal alignment is such that afferents from Δ arrive precisely in the middle of the buffer, indicated in all panels of Figure 6 as blue circle. Then, there is ½*T*_Δ_ time for consolidation, and ½*T*_Δ_ time for prediction of immediate next place cells. Note that two distinct phases of retrieval and consolidation relative to Theta were previously described in the model by Hasselmo, Bodelón, et al., 2002, and was also suggested by Dragoi and Buzsáki, 2006 and Lubenov and Siapas, 2009. In particular, Hasselmo, Bodelón, et al., 2002 found that memory retrieval and consolidation was best when entorhinal inputs to CA1 were in phase with Theta.

The rate of TPP of place cells depends on the width of their place fields (Dragoi and Buzsáki, 2006; Skaggs et al., 1996). That is, place cells with larger place fields spike later relative to Theta than cells with smaller place fields. A recent study modelled this effect mathematically and found that consistency with respect to distance between place fields is only preserved when TPP depends on the firing field width (Leibold and Monsalve-Mercado, 2017). The increase of the temporal integration window of spatial transitions, presented in Subsection 3.5, is complementary to these findings and model. Specifically, the size of the temporal integration window increases from one scale to the next by 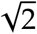. Essentially, this implies that the activation of a transition on a larger scale is slower than the activation on a smaller scale. Hence, this predicts that phase precession of a grid cell will be later with respect to Theta along the dorso-ventral axis due the scaling of the temporal integration window. More precisely, the temporal distance between spikes of individual grid cells during precession is expected to scale by 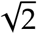 from one scale to the next.

In addition to a temporal (ring) buffer, the brain might rely on other mechanisms to support the formation of large scale transition knowledge, for instance Sharp Waves and Ripples (SPW-R) (see O’Keefe and Nadel, 1978 and review of rhythms related to Theta in Colgin, 2016). During SPW-R, actually traveled trajectories are replayed in a temporally compressed manner, which was found to support the consolidation of spatial memories (Jadhav et al., 2012). It is exactly this form of temporal compression that is necessary to form large scale transition knowledge that would otherwise not fit into the temporal integration window of STDP.

As of now, this interpretation of Theta and TPP leaves several questions with respect to TSS unanswered. For instance, it is unclear how the discontinuity between symbols in the temporal ring buffer is achieved. Possible candidates are distinct input streams arriving at opposing ends of Theta (Brankačk et al., 1993; Buzsáki et al., 1983). In addition, it was discovered that grid cells are differentially phase locked (Newman and Hasselmo, 2014). That is, there are distinct functional groups of grid cells which spike predominantly either at the trough or the peak of Theta. Conclusively, future work will address and incorporate these findings into the temporal buffer mechanism of TSS.

### 5.3 Relationship to other vector-based navigation models and successor representations

The Transition Scale-Space (TSS) is related to several other models of grid cells, and in particular to models of vector-based navigation in the rodent brain. In these models, multiple scales of grid cells are commonly used to encode a single spatial location, or a vector to a location.

For instance, the hierarchical goal directed navigation model presented in Erdem and Hasselmo, 2014 and also used in Erdem, Milford, et al., 2015 plans trajectories by first computing approximate locations using the highest scale of its representation, and narrows down on the exact position using finer scales. Their model, itself an extension of their previous work (Erdem and Hasselmo, 2012), consists of head direction, persistent spiking, grid and place cells, as well as theoretical reward cells. Because, in their model, grid cells receive head direction modulated inputs from persistent spiking cells, the model falls into the category of oscillatory interference models for grid cell formation. Moreover, it uses grid cells for localization because the activity of any place cell is defined by integrating the activity of multiple grid cells. In turn, each place cell is bi-directionally connected to a reward cell. The functional role of these theoretical cells is to provide in- and output connectivity, and represent rewards. They are a simplification of prefrontal cortical columns that were used in their previous work Erdem and Hasselmo, 2012. Their model operates in two different phases. During exploration, the first phase, an animat randomly explores an environment and recruits place cells. The recruitment is either triggered when there is currently no active place cell, or by a Poisson process that is parameterized on the inter-recruitment-interval. During the second phase, the animat first activates the reward cell that is associated with its goal location. Then, it sends linear look-ahead probes with different heading directions that start at the animat’s current location through the network of head direction, persistent spiking, grid, and place cells. When one such probe activates the representation of the goal location, the animat can navigate along the linear path towards the goal. Using just a single scale, this model has several issues. Most important, linear look-ahead probes are prone to miss the target especially when the distance between current location and target is large. The reason is that, although the angle between probes remains constant, the absolute distance between probes increases with distance from the animal. Thus, chance for missing the goal location increases. Another is that, given a neural implementation, it is to be expected that noise accumulates over time and reduces the accuracy of the probes. Finally, the authors note that reward activity might diffuse over time. Their extension of this model to multi-scale representations addresses these issues as follows (Erdem and Hasselmo, 2014). Instead of having a single scale of place cells, the recruitment process creates place cells of several scales. The scale increment is motivated to be inversely proportional to the firing field radius of place cells. Then, during the second phase, linear look-ahead probes are more likely to activate a goal place cell due to the increased firing field radius of the cell. Upon activation of a goal reward cell, the reward cell then also activates all reward cells on levels with smaller firing field radii. Intuitively, the model thus first coarsely localizes the target, and then narrows down on the exact goal.

MTS, introduced in Waniek, 2018, and TSS, have several similarities to the model presented by Erdem and Hasselmo, 2014. For instance, TSS also operate in two phases: exploration during which symbols *σ_i_* ∈ *M*_Σ_ (place cells) are recruited and associated with spatial transitions *σ_j_* ∈ *M*_Σ_ (grid cells) and temporal transitions *π_k_* ∈ *M*_Π_, and retrieval, during which the system is iterated recursively and computing potential trajectories that are outbound from an agent’s location. Likewise Erdem and Hasselmo, 2014, TSS recruit symbols either randomly or by distance to previous symbols. Note that in Section 4, the exploration phase was mostly replaced by random sampling from the input space. The descending mode of trajectory computation, presented in Subsection 4.3, is particularly similar to the approach described in Erdem and Hasselmo, 2014. Both models first locate a target location in the coarsest representation. However, whereas the model by Erdem and Hasselmo, 2014 then narrows down at the target location, the TSS model generates sub-goals in the vicinity of the agent and re-plans to these sub-goals. This is due to the basic idea of TSS to expand a path from start to target into a coherent and connected sequence of intermediate symbols (Waniek, 2018). This is fundamentally different from computing a linear target vector and, thus, from Erdem and Hasselmo, 2014 or other navigation models that are based on global vectors. Specifically, and in contrast to Erdem and Hasselmo, 2014, the information that is stored within a TSS is not a sparse representation of locations, but a connected graph in which symbols (i.e. vertices of the graph) are connected explicitly via spatial and temporal transitions (edges of the graph). In addition, TSS only require iteration through grid and place cells during retrieval. Thereby, TSS are independent of the heading direction during exploration. This is exemplified in Appendix E, which demonstrates that TSS compute the geodesic between points and not a linear vector if the input space is a non-Euclidean manifold. Note that path expansion via traversal of a connected graph also means that an TSS can compute trajectories that would require the agent to first walk into an opposing direction. For example, assume an environment with a large U-shaped barrier and in which the goal is to navigate from the center of the environment to a location behind the barrier. Then, linear look-ahead probes fail to accomplish the task, whereas TSS successfully compute trajectories. One drawback of this is that the necessary connectivity of the represented graph means that TSS cannot compute trajectories to locations that were not previously explored. However, this is in line with evidence from rodent experiments in which rats were unable to take novel shortcuts (Grieves and Dudchenko, 2013). There are several other significant differences, as for instance that grid cells are not assumed to form on the basis of oscillatory interference but because of dense packing of receptive fields that represent transitional information (Waniek, 2018). In addition, place cells were hypothesized to be recruited from spatial afferents independently from grid cells. Recent evidence from rodent experiments supports this idea (Chen et al., 2019). Note also that, although this paper uses multi-scale grid cells, it does not recruit multi-scale place cells. Another difference is the form of activity during retrieval. Whereas linear look ahead probes lead to targeted and directional activity, the search strategy of TSS resembles a propagating wave of activity from the goal to the target (see Figure 8). The latter is consistent with evidence that shows that activity in the Hippocampal formation propagates in waves (Lubenov and Siapas, 2009; Patel et al., 2012). Unlike the model by Erdem and Hasselmo, 2014, multiple scales of representation were motivated in Subsection 3.1 not by linear probes missing the goal location or by reward diffusion, but by examining the abstract runtime complexity of sequence retrievals in transition systems and, thus, wave propagation. Finally, Erdem and Hasselmo, 2014 did not provide a mathematical justification for the scale increment in their model. In contrast, the scale increment between consecutive levels of the TSS was derived analytically for biologically plausible receptive fields and with the goal to optimally construct a data structure that stores look-ahead transitions in Subsection 3.3.

Another linear look-ahead model was presented by Kubie and Fenton, 2012. First, the authors highlight the difference between navigation with a place cell map, in which a path can be computed as a trajectory that minimizes the place fields that are crossed between goal and target, and vector navigation using grid cells. The authors note that the regularity of the grid pattern should contain sufficient information to compute linear vectors from start to a goal location even across previously uncharted territory. Based on evidence from rodent experiments, the authors then introduce the concept of a rigid module. Such a module contains only grid cells with the same scale and orientation. Moreover, the authors define a tile: a region of space with only one grid bump or, in other words, the induced Voronoi cluster of each bump of a grid cell. In total, tiles of a rigid module densely cover the input space. Then, the authors proceed to define a shortest inter-bump vector, which is the shortest vector or shortest relative shift between any two bumps that could appear within one tile. The authors continue by asking if it is possible to learn predictions about subsequent bump locations within a tile during navigation. In simulations of a virtual agent, they find that grid cells in a single rigid module cannot learn predictive codes for subsequent grid bumps using Hebbian plasticity. However, they then show that rigid modules of conjunctive cells, i.e. grid cells that are modulated by head direction, in fact express strong connectivity between cells with different bump locations within one tile and thus represent the inter-bump vector. Based on these results, the authors finally speculate that learning such predictions could be used for linear vector based navigation. The model by Kubie and Fenton, 2012 is similar to TSS only to a limited extent. For instance, both models assume that grid cells compute or represent a local relationship structure. However, Kubie and Fenton, 2012 take the grid pattern as given, whereas the hexagonal arrangement of grid fields was derived mathematically as the result of optimality considerations in Waniek, 2018. Moreover, TSS do not use grid cells for localization. Although an inherent part of detecting the start of a transition is to determine a specific input space configuration, this information is not used for downstream localization in the form of place cells, but to learn a neighborhood relationship and, thus, the adjacency between place cells and not between grid cells. Most important, however, is that the model by Kubie and Fenton, 2012 learns to predict succeeding grid representations, whereas TSS learn transitions to (or predictions of) subsequent place cells. Finally, and similar to the model by Erdem and Hasselmo, 2014, the authors note that their model has limited capabilities in complex navigation tasks. To stress this point, linear vector based navigation models cannot compute non-linear solutions such as required in the U-barrier environment that was mentioned above, or in environments or input spaces where the geodesic is the desired solution.

Prior to the discovery of grid cells, a vector based navigation model of the Hippocampus was presented by Redish, Touretzky, et al., 1996. In their model, vectors are computed due to visual landmarks and the relative positions of a start and target with respect to these. Subsequent work of the authors then introduced the idea that the Hippocampus works in different two different modes (Redish and Touretzky, 1998). During exploration, the animal acquaints itself with the environment and learns spatial representations in place cells of CA3. Then, when the animal re-enters an environment, the authors suggest that it learns routes along the previously acquired representations. That is, when an animal moves from a location *a* to a location *b*, their model increases the strength of the recurrent collaterals between these two representations given the direction *a* → *b*, but not the strength of *b* → *a*. Although these asymmetric connections are conceptually identical to transitions, TSS explicitly represent temporal transitions in a memory *M*_Π_, as briefly reviewed in Subsection 1.2. The benefit of separation is that a currently active representation in place cells is not prone to be immediately overwritten or changed due to asymmetric recurrent connectivity. That is, place cells can be stored in an auto-associative memory *M*_Σ_ in which place cell activity is a local attractor and, thus, performs pattern completion, and in contrast, transitions can be stored in a hetero-associative memory to toggle switching of local attractors in the auto-associative memory (Palm, 1980; Wennekers and Palm, 2009; Willshaw et al., 1969). Pattern completion appears to be an import concept for place recognition, and it is thus unsurprising that, based on evidence for local connectivity, associative memories were previously used to model the Hippocampal formation (Graham et al., 2010; Marr, 1971; McNaughton and Morris, 1987; Treves and Rolls, 1994). However, TSS predicted a specific strong recurrent connectivity between memories *M*_Σ_ and *M*_Π_, which was recently supported by evidence from rodent experiments (Davoudi and Foster, 2019). Another difference between the work presented in this paper and the work by Redish and Touretzky, 1998 is that TSS store spatial transitions within grid cells of memory *M*_Γ_, and this paper extends the spatial transition memory to multiple scales. Multiple scales of representation were not considered in Redish and Touretzky, 1998. The authors note, however, that SPW-R should consolidate knowledge about sequences of places. They then suggest that phase precession is a result of these asymmetric connectivities during retrieval of such sequences. Also the work presented in this paper presents phase precession as a result of asymmetric information which re- or pre-plays previously acquired sequences. In contrast to Redish and Touretzky, 1998, phase precession is speculated to be a necessary mechanism that allows the formation of multi-scale grid cell representations. Additionally, SPW-Rs are believed to allow learning very large scales that would otherwise not fit into the temporal integration window of STDP learning rules. That is, because SPW-Rs contain temporally compressed sequences, a neuron is hypothesized to be able to associate with a larger number of temporally ordered elements than would normally fit into the temporal STDP integration window during uncompressed sequences. Due to the numerous similarities between and complementary elements of the TSS presented in this paper, and the model presented in Redish and Touretzky, 1998, it appears highly likely that both models can be combined.

Cueva and X.-X. Wei, 2018 and, independently, Banino et al., 2018 observed that training deep neural networks on spatial navigation tasks leads to the formation of place and grid representations. The latter in particular argue that their system learns vector-based navigation and spatial codes. However, a closer examination of their model architecture reveals that the cells which exhibit grid codes learn predictive codes for downstream spatial representations. More precisely, the recurrent network in which the grid representations emerged receives spatially modulated input and forwards information to a planning module. Hence, grid representations appear to be, in fact, an optimality result when learning from navigational tasks in recurrent networks. Moreover, their model learns to navigate to locations at which the agent was presented with a reward. Thus, the computation that is performed by their model closely resembles prior work on Successor Representation (Momennejad et al., 2016; Stachenfeld et al., 2016).

The TSS model of grid cells is also related to other transition models of grid cells, and in particular models that assume grid cells perform Principal Component Analysis (PCA) or compute a Successor Representation (SR) (Dordek et al., 2016; Momennejad et al., 2016; Stachenfeld et al., 2016). These models compute grid like responses based on place cell inputs and represent correlate information between grid cell and place cell activity in multiple scales. For instance, higher scales of SR models represent transitions to remote locations and learn this information exclusively on place cell afferents. In contrast, grid cells of the scale-space TSS model learn their representation on a suitable input space and link to place cells only due to co-activation learning. Thereby, they relay structural information from the input space to higher cortical representations, which can then be used independently of the input space during retrieval (see Waniek, 2018 for a discussion thereof). In addition, SR models propose that grid cells learn to represent the weighted sum of rewards of distant future states. That is, these models require a certain policy by which the animal navigates the environment and also that an environment and its intermediate states are sampled sufficiently often for the representation to converge. Moreover, the animal needs to learn a distinct SR representations for each specific goal. Given recent evidence that place cells are insensitive to reward information (Duvelle et al., 2019), it remains to determine if grid cells support reward storage, or in which way reward information could be available to support learning of an SR. Other recent experiments showed that the hexagonal grid representation is, in fact, locally distorted upon changes to the reward structure of an environment Boccara et al., 2019; Butler et al., 2019. In these experiments, grid fields moved closer to a certain location if this location corresponded to a singly peaked reward. On a first glance, these findings appear to be in favor of SR models. Yet, it is unclear in which way exactly this fits into these models, and, more importantly, what exactly drives the grid distortion. In contrast to PCA and SR models, TSS are independent of any policy or reward and can be recomputed easily for arbitrary targets. Thus, TSS can be used more flexibly in different environments and in particular when the observed reward modulation is not intrinsic to grid cells, but, for instance, attached to the spatial input space. Also, TSS require only limited information about traversed states and do not need dense or repeated coverage of the input, as demonstrated in the Morris water maze experiment presented in Section 4. Nevertheless, SR models have the benefit that they can compute trajectories to target locations without any additional reward mechanism, which is currently lacking from TSS. Possible future work therefore includes to merge these models.

### 5.4 Relationship to scale-spaces, data structures, graph algorithms, and robotics

The constructive procedure presented in Section 3 generates a data structure that contains transitions in multiple resolutions. Specifically, it forms a scale-space of transitions in the form of a Gaussian or a Laplacian-of-Gaussian pyramid. This kind of data structure is well known in the computer vision and signal processing communities (Lindeberg, 1994; Lindeberg, 2010; Witkin, 1983). Among other application areas, Gaussian pyramids were used to describe biologically plausible retinal and visuocortical receptive fields (Behnke and Rojas, 1998; Georgeson et al., 2007; Young, 1987). In addition, it was shown previously that a scale increment of 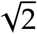 is optimal for normally distributed inputs in vision tasks (Lindeberg, 2010). In image processing, application of a Gaussian pyramid corresponds to consecutively smoothing an image, i.e. an input gets low-pass filtered by which fine-scale information is removed. Consequently, this allows to detect features in a scale-invariant way, which is commonly used in classical computer vision algorithms (Lindeberg, 2015; Lowe, 2004).

The same intuitive idea is certainly true for spatial navigation. To plan sequences to remote locations, it appears behaviorally unnecessary to factor fine details close to the target into the solution. Rather, it seems more important to quickly establish an approximate solution, which can be refined during execution. In other cases, fine details may not be relevant at all, and approximate solutions entirely sufficient. Gaussian pyramids provide a principled solution to compute these approximations, as demonstrated in the examples presented in this paper. More specifically, Figure 9 and Figure 10 show how a TSS can compute trajectories with details near the agent and approximate steps at the target, and Figure 11 show the level of approximation that each scale provides.

The computational benefits of spatial approximations with and clustering into multiple scales of representation is also well known in other areas. For instance, the computer graphics community has widely adopted data structures that are based on or extensions of quad-trees (Finkel and Bentley, 1974; Samet, 1984), or higher dimensional variants such as octrees (Orenstein, 1982), to achieve real-time graphics. Also the robotics and path planning communities have adopted multi-scale data structures (Behnke, 2004; Hwang et al., 2003; Kambhampati and Davis, 1986; Lu et al., 2011). These algorithms usually represent an environment topologically or with an imposed grid and use some variant of Dijkstra’s algorithm (W. Dijkstra, 1959) or A* (Hart et al., 1968) to determine an optimal path from start to target.

The method presented in this paper is related to these classical graph algorithms, and in particular A* (Hart et al., 1968). A* generates optimal solutions during graph searches, but its runtime crucially depends on a suitable heuristic. The basic idea of A* is the same as in Dijkstra’s algorithm, meaning that the algorithm sequentially expands nodes and tests if they are on a path to the target. In contrast to Dijkstra’s algorithm, A* uses a heuristic that guides the expansion of nodes and thereby limits the search space significantly. Still, A* requires that the heuristic is admissible, which means that the heuristic always has to underestimate the real distance to the target. The scale-space approach presented in this paper adheres to this condition, and is thus a suitable heuristic for A*.

## 6 Conclusion and outlook

In this paper, transition encoding was examined from the perspective of behavioral requirements for path planning. An example showed that a single scale of transition encoders is insufficient to provide run-times that lie in a meaningful range when neuronal axonal delays are respected. However, transition encoding allows interpretation in form of linked lists of data, for which asymptotically optimal search strategies are well known from computer science. In particular, interval skip lists introduce hierarchies to accelerate searches exponentially.

It was then derived that the optimal scale increment for a data structure with biologically plausible spatial receptive fields is 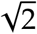. Following this, it was argued that also the temporal integration window needs to increase accordingly, and that temporal buffering of data is mandatory.j The resulting hierarchical representation, called TSS, can accelerate path planning by transition look-ahead forms a scale-space that provides approximate path planning solutions. This was shown in simulated navigation experiments using two different modes for sequence retrieval. In addition, a simulated Morris water maze experiment highlighted computation of approximate solutions on higher scales.

Given that scale-spaces are well known from other cortical regions, primarily the visual cortex, this allowed to discuss the results in terms of a general principle that underlies cortical processing. Furthermore, the TSS was related to techniques for multi-resolution path planning that are well-established in the robotics community. Conclusively, the discretized EC could be explained as an effective, yet general purpose, data structure that achieves optimal run-time behavior during goal-directed sequence planning.

Future work will include a biologically plausible model for the self-organization of the dendritic tree of grid cells and, thus, the formation of hexagonal grid fields in a dynamical model. Moreover, the TSS will be extended with reward propagation mechanisms and a probabilistic model of computation.

## Acknowledgments

The author would like to thank Christoph Richter and Jörg Conradt for discussions and feedback on the original idea of multi-transition encoding. Also, thanks to Christoph Schied for invaluable discussions on acceleration data structures and algorithms. Finally, thanks to Maike Krause for significant contributions to the structure of the manuscript.

The work presented in this paper was conceived primarily while the author was with the Neuroscientific System Theory Group (NST), Technische Universität München, Arcisstraße 21, 80333 München, Germany, and was partially funded by EU FET project GRIDMAP 600725. The author declares no competing financial interest.

## A A brief introduction to interval skip lists

This section briefly reviews those data structures that are of particular interest in the context of MTS.

Skip lists are a data structure that can be used to accelerate searching an element in linked lists of data. The acceleration is achieved by a hierarchy of *fast lanes* that jump over elements of the original list (Pugh, 1990). Searching an element in a skip list starts on the highest (or coarsest) level of the hierarchy and proceeds until an element is found that is larger than the target. Subsequently, the search drops down to the next finer level of the hierarchy until the element is recovered. Given uniformly sampled data, skip lists are dual to binary search (Dean and Jones, 2007) and thus become asymptotically optimal (Knuth, 1998). Specifically, the hierarchy of fast-lanes improves operations on equidistant data exponentially and retrievals, insertions, and deletions can be performed in *O*(log *N*) time with high probability. Due to their favorable properties, they found widespread application in various applications and are still actively researched (see for instance Bender et al., 2017 and references therein). Moreover, skip lists generalize to higher dimensions (Eppstein et al., 2005). Searching in a skip list is illustrated in Figure 13A.

Skip lists apply to MTS as follows. Observe that the information that is stored in an MTS forms linked lists of temporally and spatially ordered elements (see Figure 2D-G). Hence, searching specific elements in an MTS can be accelerated with a skip list. Moreover, transitions of an MTS densely and uniformly cover the input space, which is required for asymptotic optimality. However, only finding a certain element in accumulated knowledge is insufficient for most meaningful behavior, in particular during spatial navigation. In such a case, a path *A* ↝ *B* needs to be expanded into all intermediary symbols.

Retrieval of all intermediary items between two elements can be computed with *interval queries*. Given a suitable data structure *𝒟*, for instance an array of elements, an interval query iq(*𝒟, q_s_, q_e_*) returns all intervals in *𝒟* that are between query start *q_s_* and query end *q_e_*. Like binary search, interval queries can be performed in logarithmic time with suitable data structures, such as segment trees (de Berg et al., 1997), or interval skip lists (Hanson, 1991). Figure 13B depicts searching an element using an interval skip list. Answering iq(*𝒟, q_s_, q_e_*) is performed by first locating the interval that contains *q_s_*. Then, intermediate intervals are retrieved until the interval that contains *q_e_* is found. This is typically performed on the highest available level for performance reasons. Intermediate low-level values can be extracted by traversing the lower levels of the data structure on the chain of higher-level intervals. Thus, reporting all intermediate *K* values between *q_s_* and *q_e_* can be performed in *O*(log *N* + *K*) time. Moreover, segment trees or interval skip lists can be built bottom up in *O*(*N*) time (de Berg et al., 1997). Each additional layer in the tree or list contains nodes with intervals that enclose the nodes of the lower level, illustrated in Figure 13B. Likewise binary trees, segment trees and interval skip lists generalize to higher dimensions.

## B Convolution of Gaussian receptive fields

Following Subsection 3.3, let

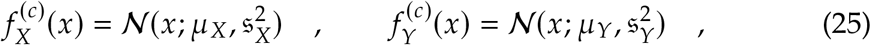

be the on-center regions of two neighboring receptive fields with preferred stimulus *µ_X_* and *µ_Y_*, respectively.

In addition, note that

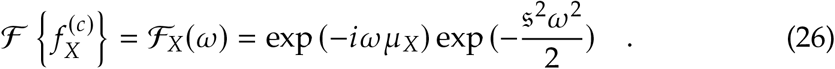

Using the convolution theorem, the on-center 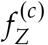, which combines 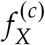 and 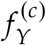, is given by

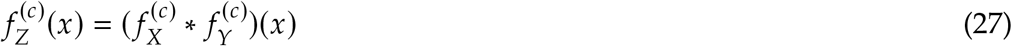

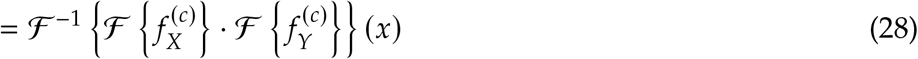

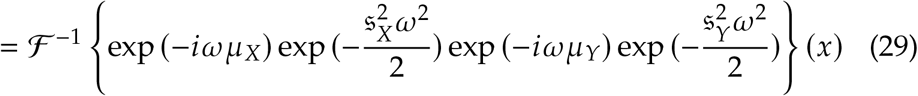

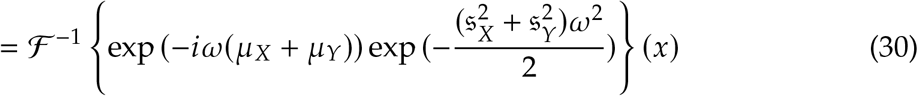

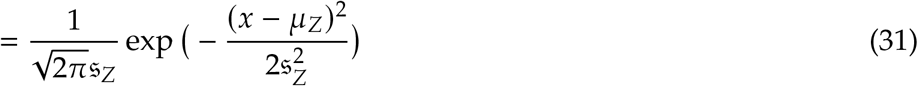

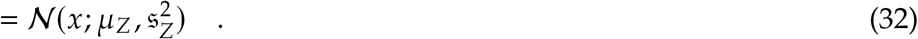

## C Algorithms

The following pseudo-code uses member-notation similar to common programming languages in the form *σ.*parents, where parents is a member of symbol *σ*. Also, multiple assignment instructions may be stated in a single line (e.g., Algorithm 4, line 2). An implementation in python of the algorithms and additional code for visualization can be found at https://github.com/rochus/transitionscalespace.

Algorithm 1 computes the expansion of a set of symbols. That is, given a set of transitions Γ and currently active symbols *S_t_* at time *t* it computes all subsequent symbols of Σ that can be reached and have not been expanded previously. The algorithm also updates the parents of all such newly activated symbols. A transition *γ* has members image and domain, which are sets containing the symbols in the image and domain of the transition.

Given a set of transitions Γ, a set of symbols Σ, a set of starting symbols Σ_start_, and a set of target symbols Σ_target_, Algorithm 2 computes a sequence from any of the start symbols to any of the target symbols on one scale of transitions. The algorithm does so by calling Algorithm 1 repeatedly until the target symbol is in the set of currently active symbols.

Backtracking a solution sequence from target to start is computed as specified in Algorithm 3. That is, given a set of symbols Σ, a set of starting symbols Σ_start_ and a set of target symbols Σ_target_, a path is computed by repeatedly randomly selecting a parent node until one starting symbol is found. Note that this requires that Algorithm 1 was applied to the symbols, whereby the parent field of each symbol is set accordingly.

### Algorithm 1 Expanding a graph to all subsequent symbols

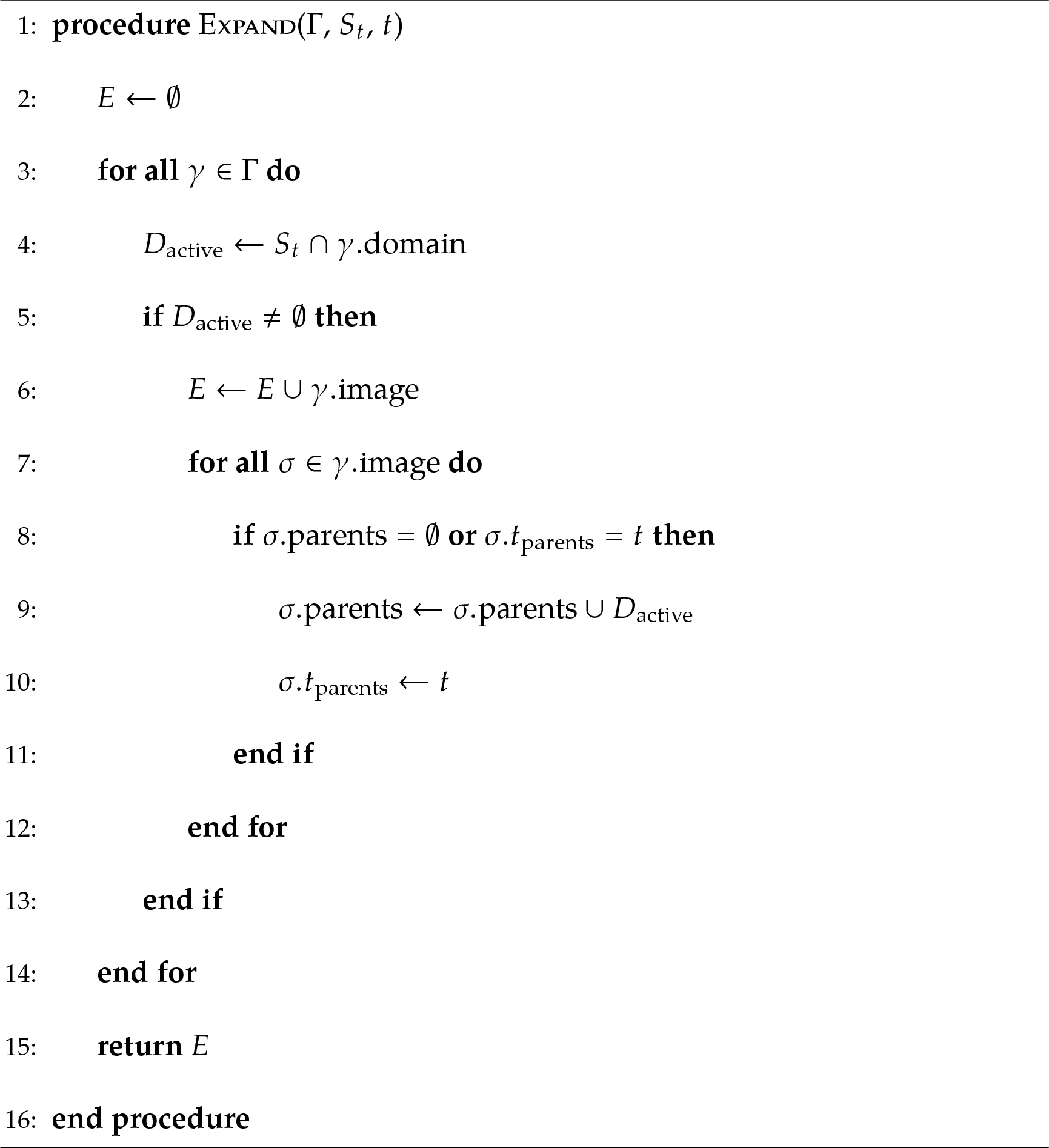

### Algorithm 2 Find a sequence on one scale of transitions with flood fill

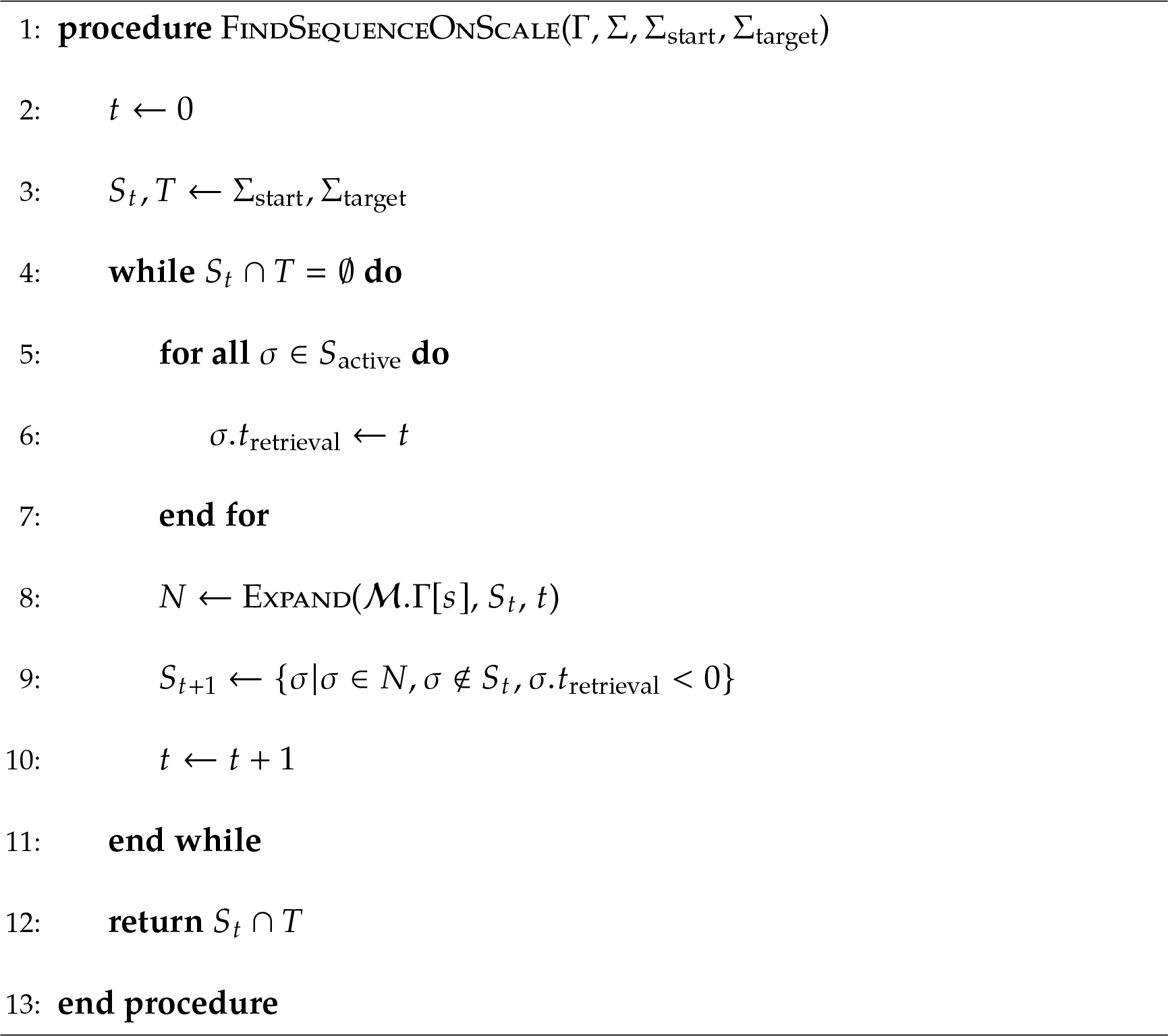

### Algorithm 3 Backtrack from target to start

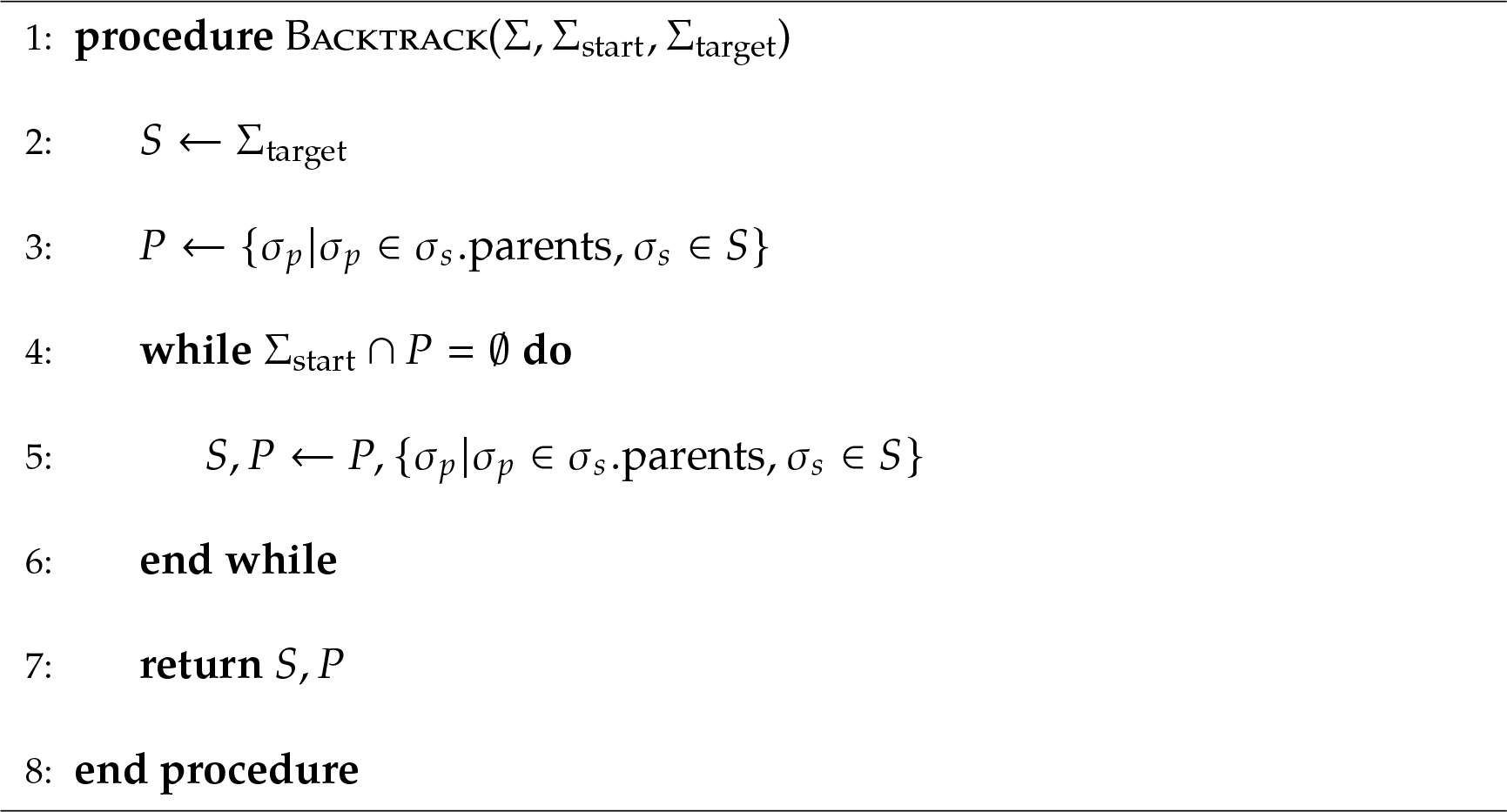

Algorithm 4 takes a scale-space MTS *ℳ*, a set of symbols Σ, a set of start symbols Σ_start_, a set of target symbols Σ_target_, and a number of maximal expansions per scale. It then computes at most *i*_max_ expansion according to Algorithm 1 per scale. If this maximum number is reached, the algorithm escalates to the next scale. *ℳ*.Γ[*s*] denotes transition scale *s* of *ℳ*. Afterwards, Algorithm 3 can be used to compute a Monte Carlo solution sample.

Algorithm 5 takes the same arguments as Algorithm 4 except the maximal number of scales. The algorithm starts on the highest available transition scale and finds a sequence with it. After finding a solution, it backtracks to the starting symbols and generates sub-goals. Subsequently, it drops down one scale to refine the trajectory to the sub-goals. In its current form, this requires to reset the parent information in each symbol (line 6).

### Algorithm 4 Find a sequence of symbols by ascending the scale-space

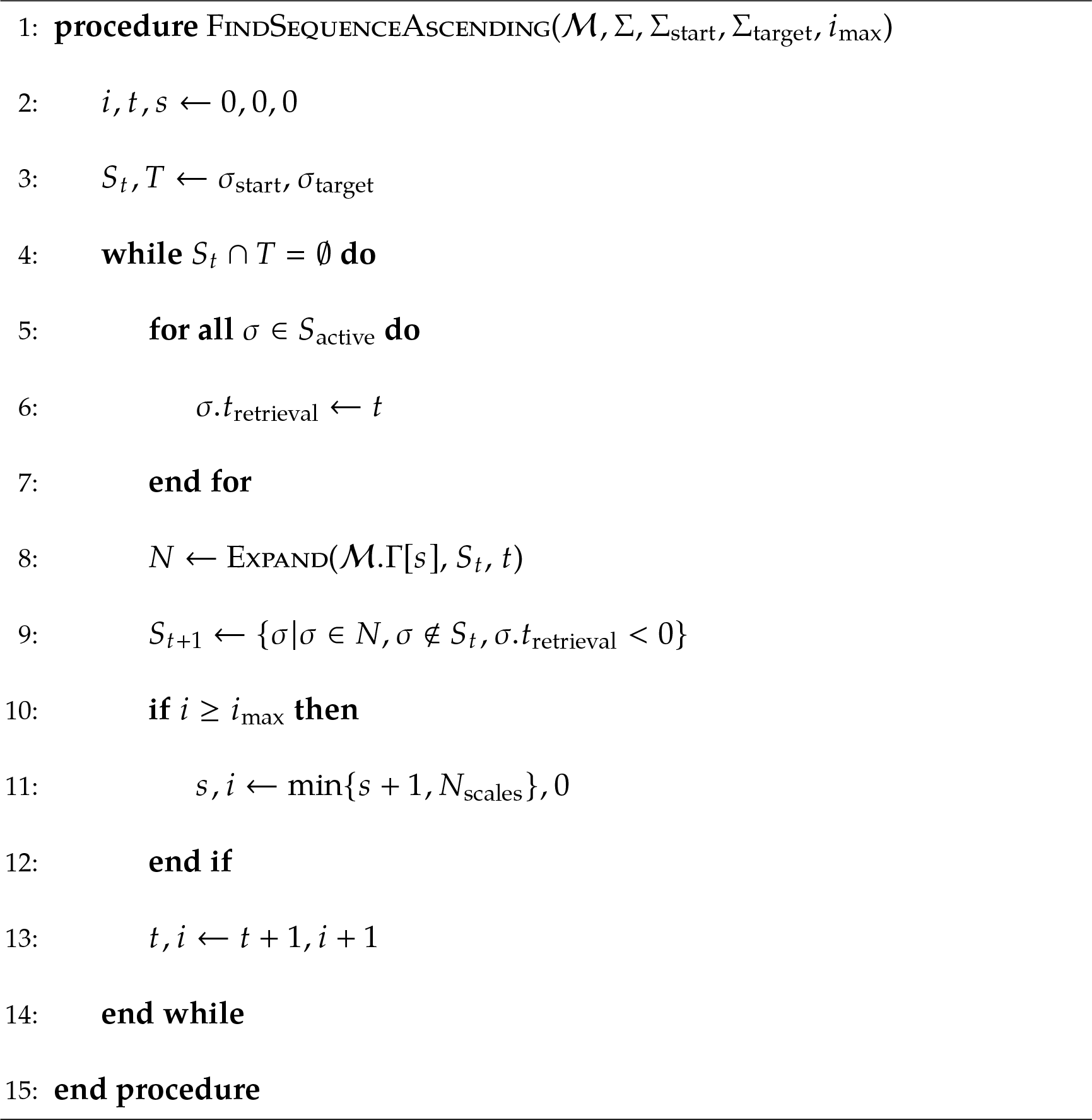

### Algorithm 5 Find a sequence of symbols by descending the scale-space

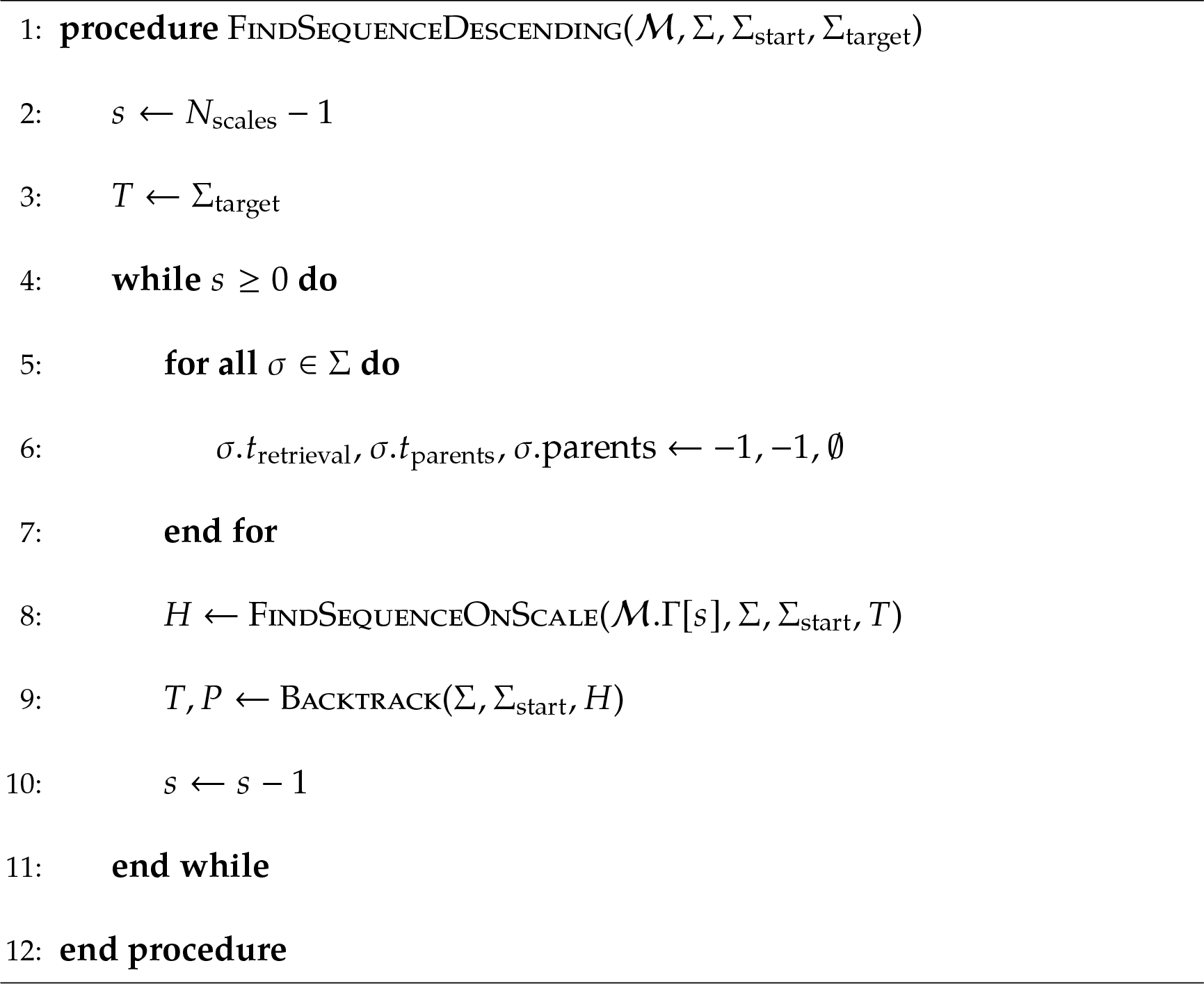

## D Relationship to category theory

Multi-Transition System (MTS) as proposed in Waniek, 2018 and extended to a scale-space representation in this paper have a relationship to category theory (see Adamek et al., 2009 for an introductory text). Changing the terms that were used throughout this paper makes this connection more pronounced. Let any transition between two symbols of the same alphabet be a *morphism* that operates on some objects. For instance, spatial transitions are morphisms *g* : *x* → *y* that operate on objects *x, y* of the category *I* of spatial symbols. Learning spatial transitions, part of the proposed dendritic computation of grid cells, is thus learning of morphisms in the category *I*. In MTS however, grid cells simultaneously also learn transitions in the representation that is created by place cells, i.e., morphisms *f* : *p* → *q* that operate on the category of place symbols *P*. Observe that, according to scale-space MTS – but also in the context of Successor Representations – grid cells learn to preserve the structure of the input space and convey this knowledge to place cell representations. Consequently, grid cells can be interpreted as *functors F* that transport objects *x* ∈ *I* to objects *Fx* ∈ *P* and morphisms from the category of spatial symbols *I* to morphisms in the category of place cells *P*. Namely, they transport transitions between symbols from one category to the other, that is *F g* : *F x* → *F y*, while preserving structure. Also, for every functor *F*, there exists an inverse functor *F*^−1^, i.e. *F*^−1^ *f* : *F*^−1^ *p* → *F*^−1^ *q*. Note that these observations apply across different scales of transition representations, regardless of the size of the grid field, and thus expose a fundamental cortical operation. Figure 14 illustrates the relationship between MTS and category theory. During spatial navigation, the existence of the inverse functor allows to perform inference on the sensory inputs *δ_i_* ∈ Δ given a sequence of symbols *σ_k_* ∈ Σ. This is demonstrated by computing the solution space in the examples of Section 4. Certainly, also other cortical areas have to learn relationships between objects of different categories while preserving structure, for instance to link the visual representation of an “apple” with the word “apple”. Conclusively, it appears reasonable to investigate if cortical computations, and in particular those that learn to represent relational information, can be understood and expressed compactly in terms of category theory.

**Figure 13:**
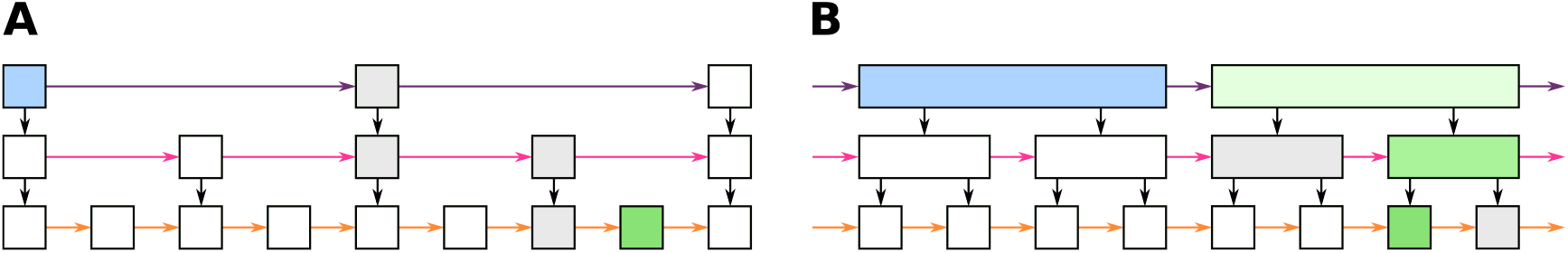
(A) A skip list is a multi-layer linked list data structure (Pugh, 1990). Elements of the list must be ordered. Higher layers have fast lanes to skip over elements. Given a start (blue), searching an element (green) in a skip list starts in the highest layer. Skip links are followed until an element on the corresponding layer is found which is larger than the target. Then, the search is continued on a lower level (see gray boxes). (B) In contrast to skip lists, interval skip lists (Hanson, 1991) store intervals in higher layers. Searching an element is similar to skip lists, but tests if the target element is in an interval (indicated by larger boxes), and subsequently drops down in the hierarchy. Interval skip lists are related to segment trees (see de Berg et al., 1997).

**Figure 14:**
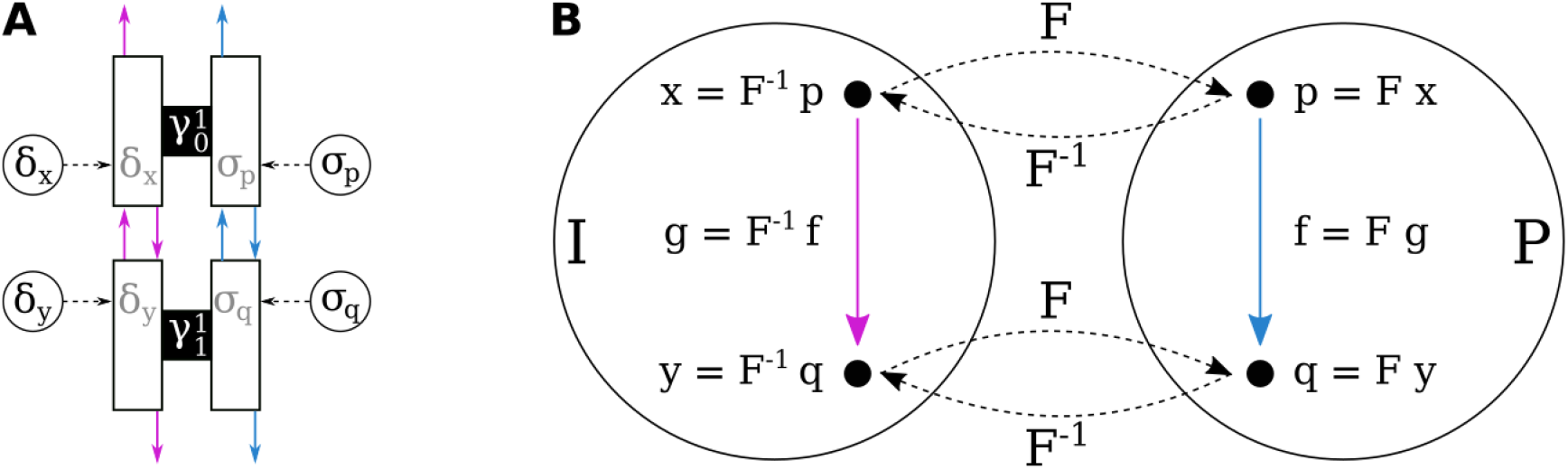
Transitions, grid cells, and category theory. (A) Visualization of grid cells according to scale-space MTS that learn transitions from symbols *δ_x_* to *δ _y_*, while also learning transitions from *σ_p_* to *σ_q_*. (B) Commutative diagram of category theory, in which a morphism *g* maps objects *x* to *y* in a category *I*, a morphism *f* maps objects *p* to *q* in a category *P*. The morphisms and objects are linked by a functor *F* that preserves structure of the categories.

## E Relationship to differential geometry

Multi-Transition Systems (MTS) hypothesize that grid cells learn spatial transitions (Waniek, 2018). Specifically, grid cells are believed to densely pack on-center and off-surround receptive fields, their grid fields, on a suitable input space. For analytical treatment, the input space was assumed to be the Euclidean. However, MTS generalize to Riemannian manifolds and are thus related to the work by Gustafson and Daw, 2011.

Gustafson and Daw, 2011 proposed that grid and place cells are modulated by geodesic distance. Their work was motivated by the idea that similarity between places should be defined locally along a path and along obstacles, and not globally due to coordinates. Using a virtual RL agent, they showed that Euclidean distance measures can have detrimental effects on learning a value function. More precisely, their model learns approximate values of locations by linearly combining a set of basis functions. In their model, these basis functions were pre-determined according to an oscillatory interference model of grid cells, i.e. as the sum of three cosine waves. The authors concluded that spatial representations in the brain should follow geodesic distances instead of Euclidean. In addition, they modelled how value functions should deform under geodesic considerations and made several testable predictions about the deformations of the grid code. Recently, experiments by two different labs showed that the grid response indeed deforms due to environmental influences (Boccara et al., 2019; Butler et al., 2019).

Likewise the model by Gustafson and Daw, 2011, MTS were motivated with the idea that navigation needs to learn similarity or relations between locally adjacent places and not across global distances. In contrast to the work by Gustafson and Daw, 2011, triangulation of the input space (i.e. hexagonal placement of grid cells) was not predefined but derived as an optimality result in Waniek, 2018. Moreover, Waniek, 2018 hypothesized that this should emerge due to local interactions of receptive fields, and showed this using simulations of a particle system with local push-pull dynamics in an Euclidean input space.

Suppose now that the input space is not Euclidean, but a Riemannian manifold. Still, assume that grid cells densely pack on-center and off-surround receptive fields on this manifold. Due to their limited extent, receptive fields and, consequently the cells to which they belong, interact only locally. Then, it appears that receptive fields capture a locally Euclidean region of the manifold, and that the crossover from one receptive field to the next is smooth. In other words, the on-center component of a single receptive field forms a chart (or coordinate system) on the manifold, and the entirety of on-centers of all receptive fields form an atlas of the manifold.

Having an atlas of a manifold allows to compute geodesics between arbitrary points. That is, it is possible to compute the shortest path between two points on a curved surface. Intuitively, this can be related to grid cells as follows. Densely packing grid fields onto the manifold leads to a surface triangulation of the manifold. Computing shortest paths on such a mesh is a well studied problem. It can be solved approximately for instance with Dijkstra’s algorithm, the Fast Marching Method (Sethian, 1996), or the Heat Method (Crane et al., 2013). The latter two techniques have in common that they directly solve a wave propagation problem, and also Dijkstra’s algorithm propagates a wave from source to target (see experiments in Section 4).

The following simulation of a virtual agent and an MTS was used to explicitly demonstrate this relationship.

The agent moved at constant speed, and its heading direction was drawn from a Laplacian distribution at each time-step of the simulation. The input space, i.e. the space on which the animal moved, was the outside of an upper hemisphere. Note that, due to symmetry, the presented results are identical to an agent walking on the inside of a lower hemisphere.

First, receptive fields and their interactions were simulated as a particle system with push-pull dynamics similar to the method described in Waniek, 2018. That is, only particles close to the agent were updated. Specifically, the closest particle to the agent was pulled closer to the agent. All other particles in the surrounding of the agent exerted relative forces on each other. The update rate with which these forces acted slowly decayed over time. The agent and particle system was simulated for an arbitrary time until the particles converged to an approximately stable solution (see 3D view in Figure 15). The figure shows that the placement of receptive fields forms a blue noise distribution on the input manifold, or a low-discrepancy sequence. Holes in the arrangement are probably due to non-uniform sampling of the input space, and because of the simplicity of the particle system. However, visual inspection (without computing gridness scores) shows that fields form a dense packing. The particle locations defined the centers of spatial transitions.

**Figure 15:**
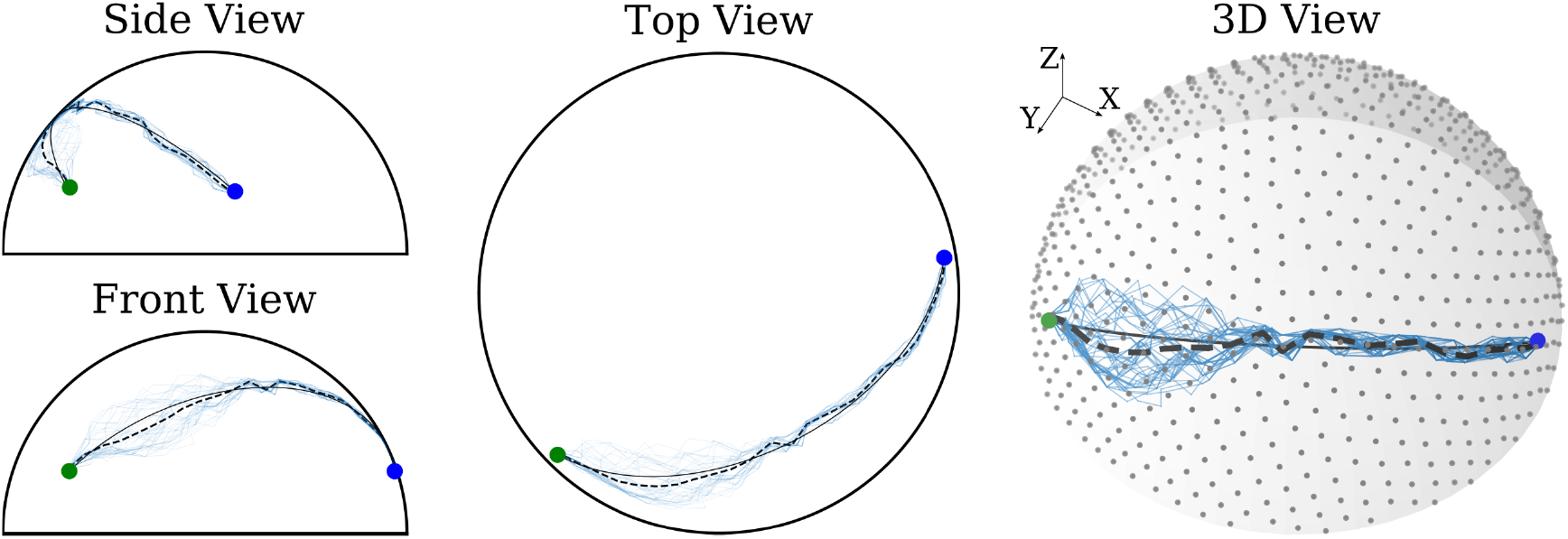
MTS on hemisphere. A virtual agent equipped with a push-pull particle system was simulated for an arbitrary time until the particle system converged. The particle locations were then used as transition centers (gray hexagonally arranged dots in 3D view). Subsequently 5000 symbols were randomly placed on the hemisphere (data not shown), and transitions and symbols associated. This formed the data contained in the MTS Then, the MTS was queried to expand a path from start (green dot) to target (blue dot). Finally, 50 Monte Carlo samples were drawn from the solution space (thin blue lines). The figures display the ground truth geodesic between start and target (thin black line), as well as the average of all Monte Carlo samples (dashed thick line). See Waniek, 2018 for description of the particle system, and Section 4 and Appendix C for details about the algorithms.

Then, 5000 points on the hemisphere were randomly sampled to be used as symbols. Subsequently, each symbol was associated with the closest spatial transition, which in turn was associated to point to all surrounding symbols. This forms the MTS.

Finally, the shortest path between two arbitrarily chosen points was computed by expansion of the MTS and backtracking 50 Monte Carlo samples (see Section 4 or Appendix C for details). The results are presented in Figure 15. Although the simulation did not yield the perfect solution, the samples and, in particular, the sample mean are close to the geodesic between the two points. The deviation from the geodesic is primarily due to the placement of the particles and distribution of symbols, and the expansion from the start symbol towards the goal symbol. That is, the MTS is not aware of real physical distance between locations, and treats all transitions with equal cost. Moreover, the search from start towards the to goal propagates in discrete steps in this simulation and thus has multiple solutions with equal cost, especially in the beginning.

Note that the presented simulation results do not imply that grid fields necessarily arrange in the depicted way when a rodent moves on a small hemisphere. Rather, the purpose of the simulation is to demonstrate that MTS can compute (approximately) shortest paths even when the input space is non-Euclidean. This appears to be particularly important given recent evidence that shows grid field deformations due to changes in the environmental reward structure (Boccara et al., 2019; Butler et al., 2019).

Source code for all simulations presented in this paper can be found at https://github.com/rochus/transitionscalespace.

## Footnotes

1 https://github.com/rochus/transitionscalespace

2 This is well reflected in the implementation that was used to simulate the TSS. It uses a single method, parameterized only by scale, for data construction. Source code available online at https://github.com/rochus/transitionscalespace.

